# Accelerated simulations of RNA clustering: a systematic study of repeat sequences

**DOI:** 10.1101/2023.12.23.573204

**Authors:** Dilimulati Aierken, Jerelle A. Joseph

## Abstract

Under certain conditions, RNA repeat sequences phase separate yielding protein-free biomolecular condensates. Importantly, RNA repeat sequences have also been implicated in neurological disorders, such as Huntington’s Disease. Thus, mapping repeat sequences to their phase behavior, functions, and dysfunctions is an active area of research. However, despite several advances, it remains challenging to characterize the RNA phase behavior at submolecular resolution. Here, we have implemented a residue-resolution coarse-grained model in LAMMPS – that incorporates both RNA sequence and structure – to study the clustering propensities of protein-free RNA systems. Importantly, we achieve multifold speedup in the simulation time compared to previous work. Leveraging this efficiency, we study the clustering propensity of all 20 non-redundant trinucleotide repeat sequences. Our results align with findings from experiments, emphasizing that canonical base pairing and G-U wobble pairs play a dominant role in regulating cluster formation of RNA repeat sequences. Strikingly, we find strong entropic contributions to the stability and composition of RNA clusters, which is demonstrated for single-component RNA systems, as well as binary mixtures of trinucleotide repeats. Additionally, we investigate clustering behaviors of trinucleotide (odd) repeats and their quadranucleotide (even) counterparts. We observe that odd repeats exhibit stronger clustering tendencies, attributed to the presence of consecutive base pairs in their sequences that are disrupted in even repeat sequences. Altogether, our work extends the set of computational tools for probing RNA cluster formation at submolecular resolution and uncovers physicochemical principles that govern the stability and composition of resulting clusters.

---

RNA serves as a ubiquitous component of biomolecular condensates [1–5] – membraneless compartments within living cells formed through phase separation. Moreover, studies have demonstrated that protein-free RNA systems can undergo phase separation [6–8]. Notably, RNA-driven condensates play crucial roles in both normal cellular function and dysfunction [2, 9, 10]. Particularly intriguing are RNA-based condensates comprised of tandem repeat RNA sequences, which have been associated with various neurological and neuromuscular disorders [6, 11]. Consequently, there exists an urgent need to decode how RNA sequence, structure, and dynamics modulate the material properties and functions of biomolecular condensates.

Recent advancements in experimental techniques, including smFISH [12], SHAPE [13], and proximity labeling [14], have offered valuable insights into the roles of RNA polymers in biomolecular condensates. However, investigating the effects of RNA at submolecular resolutions remains a formidable challenge. Computer-based approaches, particularly explicit chain molecular dynamics simulations [15–19], have proven to be excellent complements to these experimental methods, striving to bridge the gap in our ability to map RNA sequences to their phase behaviors.

In general, faithfully representing RNA in molecular simulations requires accounting for several intrinsic and extrinsic factors, including RNA charge (electrostatic effects), base stacking, base-pairing interactions, and effects due to solvent and ions (including divalent ions). However, incorporating more details is often associated with greater computational cost, especially when studying collective behaviors like phase separation. In pursuit of greater efficiency, it becomes necessary to sacrifice certain details. The resulting models are often tailored to probe specific questions of interest. For instance, it has become common practice to represent RNA at the nucleotide resolution or lower, with intramolecular interactions captured by pseudohard-sphere potentials or charged beads governed by Coulombic potentials with Debye–Hückel screening. Such models are frequently interfaced with coarsegrained protein models to investigate how the presence of RNA influences protein condensation [18, 20–22]. To enhance the representation of RNA polymers in particle-based simulations of condensates, explicit considerations for base-pairing propensities [15, 18, 19] have also been included. Notably, in the aforementioned studies, RNA has negligible propensity for secondary structure formation—i.e., the polymers behave as fully flexible chains. Significantly, for phase-separating systems, representing such features at nucleotide-resolution or higher is desirable, as it will allow one to dynamically probe how various RNA motifs influence condensation while preserving the underlying RNA sequence and may lead to emergent properties that are inaccessible via more simplistic models [23]. From a technical standpoint, capturing RNA structure has been a longstanding challenge – beyond the condensate field – and indeed, there have been several computational approaches of varying resolutions designed to address this [24]. However, in the context of simulations of condensates, where one has to model several copies of often long RNAs, accounting for secondary structure formation can come at significant computational cost due to the increase in complexity of the underlying potential energy function. In this work, starting from an existing nucleotide-resolution model of RNA that incorporates secondary structure, we question: (1) how efficient is such an approach for probing phase behavior of RNA? (2) how can we boost the efficiency of molecular simulations of RNA condensation when modeled via many-body potentials at nucleotide resolution? (3) what insights can such simulations reveal about RNA sequences and their phase behavior?

Concretely, our work builds upon the existing singleinteraction-site (SIS) RNA model [15], proposed by Nguyen and coworkers. In this RNA model, each nucleotide is modeled via a unique bead. Notably, the model parameters are derived through a top-down coarse-graining approach leveraging experimental structures of RNA repeat sequences [15]. In so doing, the model includes many-body potentials (angular and dihedral terms) to recapitulate the A-form helix of CAG-repeat sequences. It is important to note that the SIS RNA model [15] does not explicitly account for RNA charges – instead, these effects are implicitly included in the model interaction terms. Consequently, this model is most suitable for probing protein-free RNA systems in high salt regimes. Additionally, nucleotide–nucleotide interactions are scaled to account for canonical and wobble base-pairing propensities. Interestingly, this model successfully reproduces the length-dependent clustering behavior observed experimentally in repeat RNA sequences. However, more sophisticated models like the SIS model often come with an increased computational cost. As an illustration, the initial implementation of the SIS RNA model, the authors report requiring approximately 100 days of sampling to acquire statistical data on the clustering propensities of RNA. The current challenge, therefore, lies in finding a balance: how can we effectively model RNA clustering behaviors with both experimental accuracy and computational efficiency, particularly when utilizing many-body potentials?

Here, we broaden the accessibility and boost the efficiency of a nucleotide-resolution many-body potential designed for probing the clustering propensities of RNA. Our starting point is the SIS RNA model [15], initially implemented in OpenMM [25], which inherently employs ‘implicit’ force fields – eliminating the need for explicit definitions of analytical forces by practitioners. Notably, we extend the approach to LAMMPS [26], a widely utilized scientific molecular simulation software, with exceptional parallelization capabilities. To achieve this, we explicitly derive the analytical forces for the many-body base-pair potential using the formalism outlined in Ref. [27]. We seamlessly integrate these forces into LAMMPS, implementing optimizations to reduce redundant force computations and leverage parallel computing. Furthermore, we derive the analytical pressure tensor for the model, facilitating simulations that involve variations in mechanical forces on the system. Thus, practitioners can utilize approaches such as the slab technique [28] to further accelerate equilibration. As a validation of successful implementation, we perform numerical integration using the one-sided Euler method [29], which reveals consistency with corresponding analytical counterparts. Crucially, we perform benchmarks for the model across both software platforms, investigating the potential advantages and disadvantages of running these simulations. Our findings demonstrate that while using a single GPU in OpenMM can deliver reasonable performance, employing parallel computing in our implementation with LAMMPS can achieve up to approximately 5-fold speedups.

Leveraging the efficiency gains of the RNA model, we investigate all 20 non-redundant trinucleotide RNA repeats (excluding homopolymers; Supplementary Fig. S4) that have also been examined in experiments [30]. Specifically, we conduct molecular dynamics simulations on RNA trinucleotide repeat sequences, which include investigations of single-component trinucleotide repeat systems, comparisons between trinucleotide and quadranucleotide repeats and studies of binary mixtures of trinucleotide repeats. In each case, we characterize the material properties (e.g., cluster size, concentration, dynamics) of the collective systems. In line with experimental findings, our simulations unveil a robust positive correlation between the propensity to form RNA clusters and the ability to establish base pairs, specifically canonical Watson–Crick and G—U wobble base pairs. Strikingly, entropic contributions to clustering propensities of such systems naturally emerge, which have been postulated via an analytical model [31]. Additionally, we capture differences in RNA clustering between odd and even repeat sequences, which arise due to variations in base-pairing patterns. Collectively, this work enhances our capability to efficiently scrutinize RNA phase behavior via many-body potentials, opening avenues for closely investigating RNA conformational ensembles inside condensates.

## The implementation of the RNA coarse-grained model in LAMMPS is consistent and stable

The single-interaction-site (SIS) model is a 1-bead-per-nucleotide coarse-grained model designed to capture the properties of RNA in a sequence-dependent fashion. Initially implemented in OpenMM, the authors reported that it takes about 3 months to collect statistics for a system composed of 64 (CAG)_47_ chains. Since our main objective is to probe RNA clusters of similar and larger sizes, we first opted to enhance the efficiency of this model by leveraging the parallelization features in the LAMMPS software [26]. OpenMM employs its own toolkit to derive forces from potentials [25]. While LAMMPS can perform derivatives numerically (e.g., from tabulated potentials), analytical potentials are much more desirable due to their boost in computational efficiency. Consequently, we derive and implement analytical forces for the SIS RNA model in LAMMPS (Methods B).

To achieve this, we first assess the functional forms of interactions governing bonds, excluded volumes, and base-pairing in the model, Fig. 1(a). Bonded interactions, encompassing bonds and angles, are modeled via harmonic potentials that are readily available in LAMMPS [26], facilitating direct implementation by assigning corresponding parameters. Similarly, the excluded volume potential in the model utilizes the soft repulsive Weeks–Chandler–Andersen potential [32], which can be implemented in LAMMPS by shifting and cutting off the standard 12–6 Lennard-Jones potential. However, there is no existing form in LAMMPS for the base-pairing interactions, constituting a 6-body potential dependent on 4 angles, 2 dihedrals, and 1 distance [Fig. 1(a), bottom panel]. To address this gap, we implement this potential as a new extra pairwise potential in LAMMPS; i.e., the forces involved in this 6-body potential are derived analytically and encoded. Our procedure is detailed in Methods B. Briefly, the underlying basepairing potential encompasses a total of 66 different force components in the 3-dimensional Cartesian coordinate system. In comparison, typical Lennard-Jones type pairwise interactions commonly employed in coarse-grained models only necessitate consideration of 6 force components. The implemented 6-body potential can now be accessed via base/rna in LAMMPS, which allows users flexibility in setting preferred distance, angle and dihedral parameters. For example, left-handed helices can be accurately modeled by primarily adjusting the dihedral term.

**FIG. 1.**
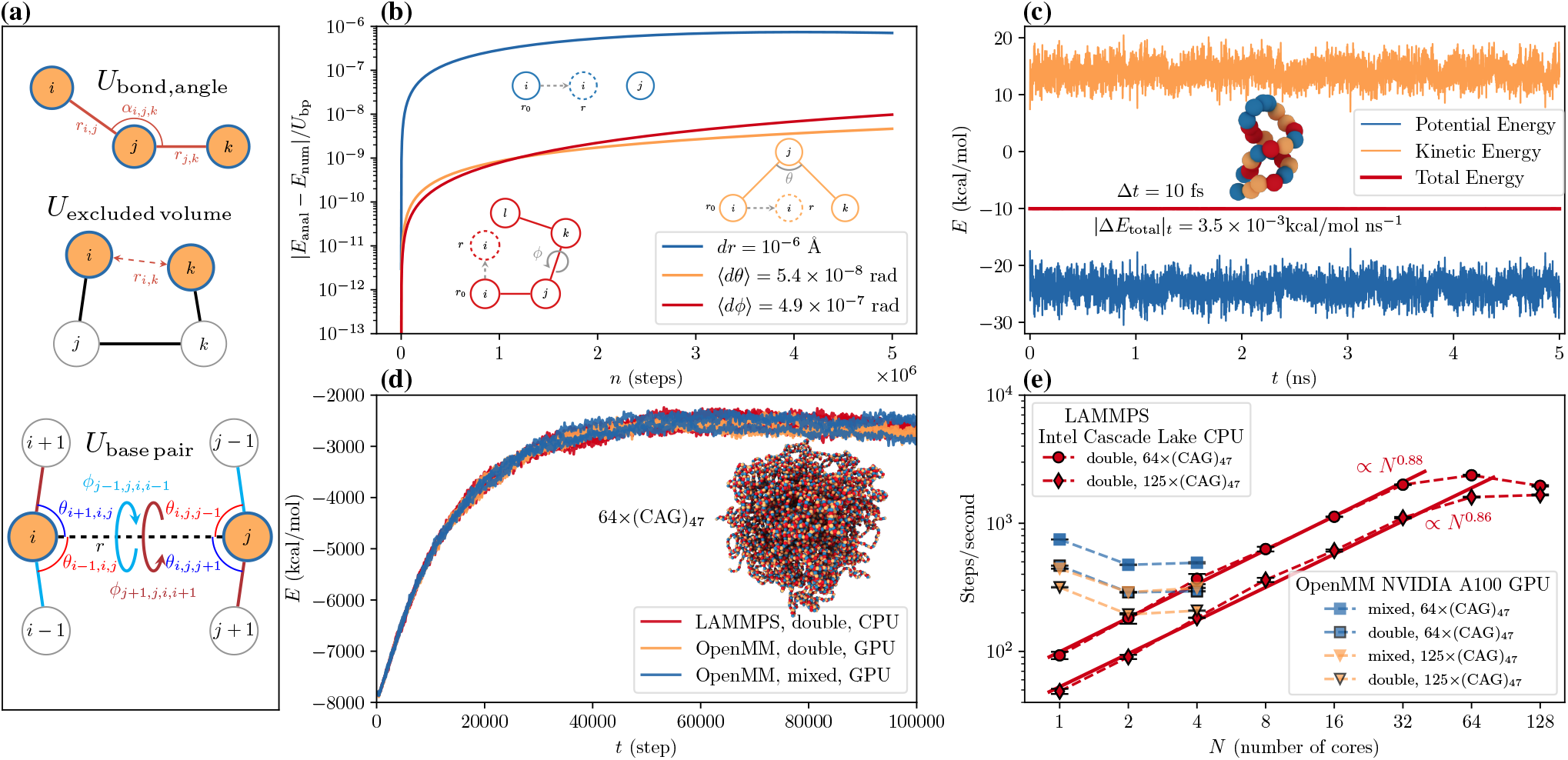
Characterization of the RNA model implementation in LAMMPS. **(a)** Schematic of potential energy terms are included in the RNA model. Namely, *U*_bond, angle_ represents the bond and angle terms along the chains; *U*_excluded volume_ captures excluded volume interactions; *U*_base pair_ represents the base-pair potential terms, which include a harmonic base-pair distance term, four angles and two dihedrals. *U*_base pair_ ensures that the A-form right-handed antiparallel helical structures are energetically favored. **(b)** Comparison between analytical and numerical integration of the base-pair potential. Numerical integration is performed via the one-sided Euler method using our LAMMPS code. We perform integration for each base-pairing term (harmonic distance, angles, and dihedrals); the specific scenarios are sketched in the plot. **(c)** Energy trajectories from the microcanonical simulation of a single chain CAG repeat. The simulation is started from a folded structure to make sure the force calculations are properly implemented. Setting the timestep size of 10 fs, the average total energy drift (3.5 *×* 10^*−*3^ kcal*/*mol ns^*−*1^) is about 4 orders of magnitude smaller than the potential energy fluctuations (*∼* 10 kcal*/*mol) demonstrating the stability of the model. **(d)** To further evaluate the correctness of the LAMMPS implementation, we simulate a condensed system of 64 (CAG)_47_ chains with Langevin thermostat at 293 K for 10^5^ steps. This compact system requires an extensive evaluation of base-pair potentials at each timestep, thus representing the slowest simulation speed. Even with different simulation softwares, different precision, and different hardware, the system follows the same relaxation dynamics. **(e)** The scaling of the implementation is evaluated for 64 (CAG)_47_ and 125 (CAG)_47_ chains. For up to 32 CPUs, the scaling exponent is 0.88 for 64 (CAG)_47_ chains. When the system size is increased to 125 (CAG)_47_ chains, the scaling exponent is 0.86 for up to 64 CPUs. Data for the original OpenMM code on A100 GPUs with both double and mixed precision are also provided for comparison.

To evaluate the correctness of our implementation, we perform numerical integration of each derived force terms for the 6-body potential [Fig. 1(a), bottom panel] via the one-sided Euler method [Fig. 1(b)]. Concretely, we test different scenarios, encompassing variation in pair distances, angles and dihedrals. Moreover, we directly compare our results to the analytical analogues, which reveals consistency for each term with respect to the integration step sizes, Fig. 1(b). Next, to assess the stability of our implementation, we conduct simulations in the microcanonical ensemble, Fig. 1(c). In these simulations, total energy is conserved while potential energy and kinetic energy undergo interconversion. The correctness of force implementation and the handling of critical terms can be further evaluated by observing the drift in total energy over time. Specifically, we prepare a folded RNA chain, ensuring extensive evaluation of base-pair interactions. As illustrated in Fig. 1(c), with a discrete timestep of 10 fs, the energy drift per nanosecond (equivalent to 10^5^ timesteps) is four orders of magnitude smaller than the potential energy fluctuations. This observation confirms the stability of our implementation.

With the stable implementation of the force field, we proceed to replicate quantitative results for individual RNA chains. In the original model parameterization, a single chain was generated by modifying the (CAG)_2_ duplex (PDB 3NJ6)[33]. The root-mean-square distance (RMSD) from the PDB crystal structure was then determined. Accordingly, we conduct simulations of the same chain and assess the RMSD, Supplementary Fig. S2. The computed average RMSD value is 5.3 Å, with a minimum RMSD of 1.1 Å. Notably, these RMSD values closely align with previously reported findings, specifically an average of 5.4 Å, and a minimum of approximately 1 Å [15]. These results further bolster confidence in the correctness of our LAMMPS implementation, particularly in capturing the structural properties of individual RNA chains.

As a final assessment of the consistency of our implementation, we perform simulations of 64-chain CAG-repeat systems. These simulations test the ability of the implementation to capture collective behaviors of RNA. Notably, we first perform an *NPT* simulation at high pressure to accelerate the condensation of the chains – which is enabled by our implementation of pressure tensors for this many-body potential (see Methods B). Starting from the same compressed structure, we conduct an *NV T* equilibration in both OpenMM and LAMMPS, Fig. 1(d), with different hardware and precision (mixed and double). The resulting trajectories are all in agreement with each other Fig. 1(d). Finally, we assess the model’s ability to replicate the clustering propensities of CAG repeat sequences. Experimental evidence indicates that such sequences display length and concentration-dependent phase separation behaviors [6]. For our evaluation, we choose two systems: (CAG)_20_ at 50 *µ*M and (CAG)_47_ at 200 *µ*M, which are expected to be in the dilute and condense regimes, respectively [6]. Consistent with both experimental observations [6] and prior simulations [15], we quantitatively reproduce the clustering propensities of these two systems (Supplementary Fig. S3). Furthermore, our computed concentrations of the clusters align well with the reported distribution of cluster concentration previously documented for (CAG)_47_ at 200 *µ*M [15].

Altogether, these analyses reveal that our implementation of the RNA model in LAMMPS is robust both in terms of capturing single- and multi-chain behaviors of RNA repeat sequences.

## Our LAMMPS implementation is scalable and multifold speedup can be achieved

The scalability of our implementation is a crucial factor for the broader applicability of the RNA model. While GPU usage has seen significant growth in recent years due to their efficiency over CPUs, CPUs remain more widely accessible on laptops, desktops, research clusters, and supercomputers. Additionally, LAMMPS is designed to be highly parallel, employing advanced domain decomposition techniques [34]. To evaluate the scalability of our implementation, we simulate condensed systems of (CAG)_47_ with 64 and 125 chains, respectively. In the case of condensed systems, numerous force calculations are required, representing the slowest simulation speed. Using the same initial conditions, we vary the number of CPUs from 1 to 128 [Fig. 1(e)]. Up to 32 and 64 CPUs, the speedup scales well in the linear regime (with a scaling exponent of *∼*0.88 and *∼*0.86) for 64 and 125 chains, respectively. Unsurprisingly, when the number of CPUs equals or exceeds the number of chains in the system, the domain decomposition and inter-CPU communication increase, leading to a reduction in simulation speed [Fig. 1(e)]. Overall, these results demonstrate that the scalability holds well as the system size increases.

In addition, we compare the LAMMPS simulation efficiency with that of OpenMM. Notably, in previous work the OpenMM simulations were performed on a single NVIDIA Quadro RTX 5000 graphics card. Since we did have access to this hardware, we use NVIDIA A100 Tensor Core 80GB graphics cards. We find that for a 64-chain system, OpenMM simulations with mixed precision running on one GPU completes 747 steps/second and going from mixed precision to double precision reduces the simulation speed for OpenMM using A100 GPUs. For comparison, the maximum speed of GPU (mixed precision; OpenMM) is equivalent to 10 CPUs in our LAMMPS implementation (double precision), rendering the OpenMM implementation advantageous in this aspect. Interestingly, increasing the number of A100 GPUs leads to negative scaling for the OpenMM implementation. This behavior could be attributed to several factors regarding the multi-GPU implementation: (1) how the GPUs are connected to each other (NVLink or PCIe); (2) the fact that OpenMM employs force evaluation decomposition instead of domain decomposition [35]. In contrast, our LAMMPS implementation achieves a maximum speed of 2370 steps/second for the 64-chain system when utilizing 64 CPUs – i.e., a roughly 3-fold increase in speed compared to the tested OpenMM GPU (mixed precision). In context, this result means that a 3 week long simulation in the original code (run on A100 GPUs) should now take 7 days to complete in the LAMMPS implementation on CPUs. It is important to note that the A100 GPUs are better suited for deep learning computations [36] and the new RTX4070/80/90 cards could yield better performance for molecular dynamics simulations [37]. Thus, the speedup in OpenMM could improve if one has access to these GPUs. Collectively, our comparisons reveal that while the OpenMM implementation holds advantages over multiple CPUs, the maximum speed of the model in OpenMM is capped in our test cases. Conversely, our LAMMPS implementation demonstrates the potential for multifold speedup – given enough resources.

## Energetic and entropic effects of base-pairing regulate material properties of RNA trinucleotide repeat clusters

Capitalizing on the improved efficiency of the RNA implementation in LAMMPS, we employ the model to comprehensively investigate the clustering propensities of RNA trinucleotide repeats (Fig. 2). RNA clusters represent regions with elevated local RNA concentrations, which may or may not result from phase separation. Within this framework, an RNA condensate denotes a cluster formed through the phase separation of RNA chains, characterized by a well-defined interface. In our simulations, we focus on RNA clusters, in general, and define these using a chain-to-chain distance metric; i.e., nucleotides belonging to chains that are within 120% of the ideal base-pairing distance are assigned to the same cluster. Using this metric, we quantify, under the same conditions, the clustering propensities – the cluster size, concentration, and the fraction of chains in the largest cluster – of various protein-free RNA systems.

**FIG. 2.**
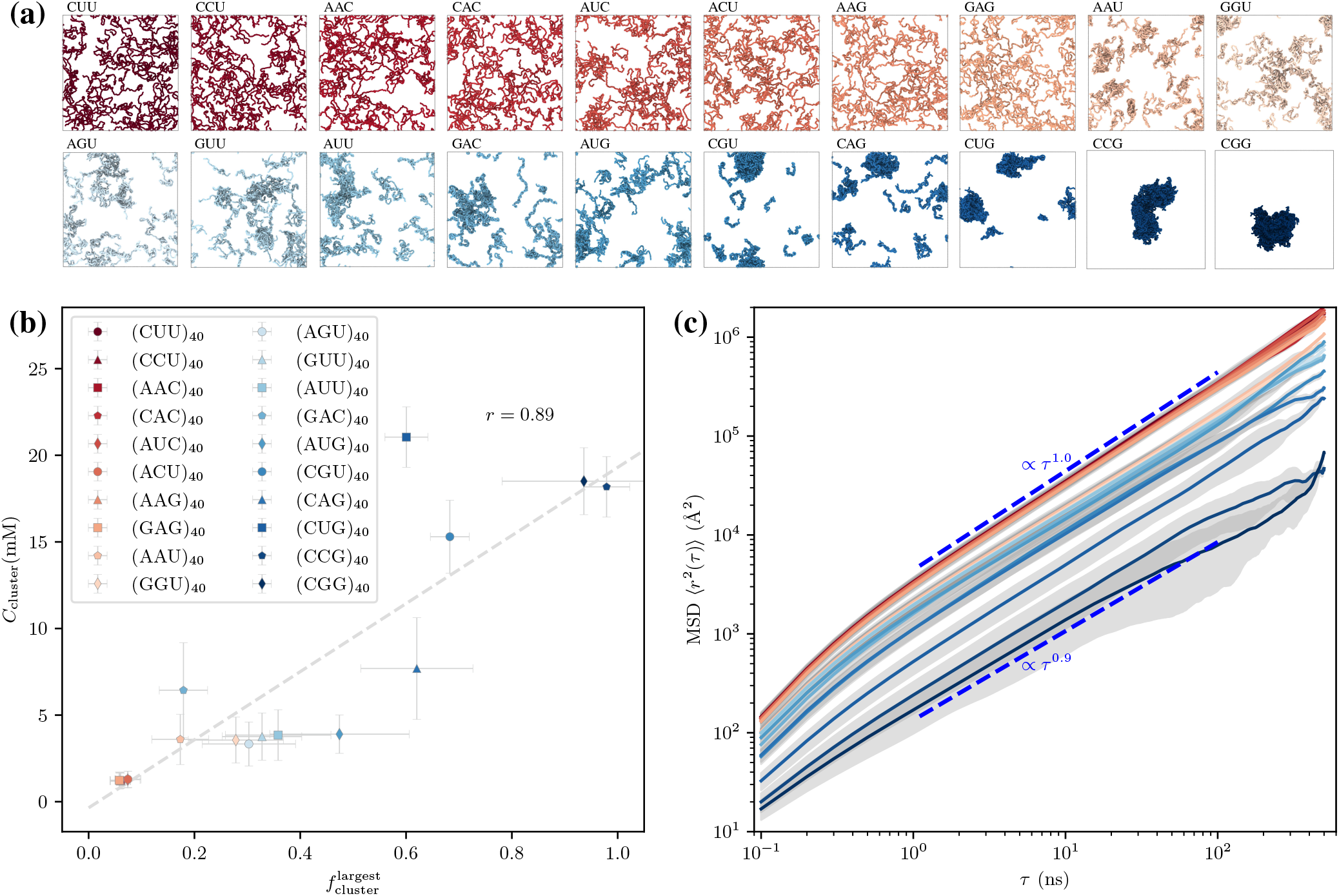
Material properties of non-redundant trinucleotide RNA repeat clusters. Each simulation is prepared by placing 64 chains of 40 trinucleotide repeats evenly in a periodic box. After using an annealing procedure (described in the text), we simulated each system for 2 *µ*s in the *NV T* -ensemble at 293 K. **(a)** Depicted snapshots are taken from the end of the molecular dynamics simulations. Corresponding sequences are labeled at the left top of each snapshot. The redundancy of the sequences is explained more explicitly in Supporting Fig. S4. **(b)** Fraction of chains in the largest cluster versus concentration of the largest clusters, indicating strong positive correlation between clustering propensities and the concentration of the resulting clusters. Here, two chains are assigned to the same cluster if at least two of their nucleotides are within 1.2 *r*_0_, where *r*_0_ is equilibrium base-pairing distance. Error bars represent standard deviation. **(c)** Mean squared deviation (MSD) over time *τ* of chain center of mass. MSD exponents of select trajectories are indicated on the plot. Error bands represent standard deviation.

Recently, it has been demonstrated that protein-free RNA systems undergo lower critical solution temperature (LCST) phase transitions [8]. Furthermore, in the phase-separated regime, at even higher temperatures, such systems can undergo a percolation transition, sustained by intermolecular base pairing and base stacking interactions [8]. Interestingly, RNA LCST phase transitions can lead to hysteresis where, upon lowering the temperature, percolated clusters still persist. Moreover, for such systems outside of the phase separated regime (sub-saturated solutions) an assortment of clusters will be formed [20, 38]. Hence, the clustering propensities we measure provide insights into the ability of given RNA chains to form droplet-spanning networks.

Specifically, we study the clustering propensities for the complete set of non-redundant trinucleotide repeats, Supplementary Fig. S4. In total, there are 64 trinucleotide repeat combinations, with the CAG repeat being just one among them. Excluding homopolymers and accounting for redundancy, this leads to a set of 20 non-redundant trinucleotide repeats, Supplementary Fig. S4. To systematically investigate the clustering propensities of the 20 non-redundant trinucleotide repeats, we adopt the similar methodology as employed for (CAG)_47_; simulating systems composed of 40 repeats at 200 *µ*M [6, 15]. Concretely, we define an RNA cluster by using a chain-to-chain distance metric; i.e., nucleotides belonging to chains that are within 120% of the ideal base-pairing distance are assigned to the same cluster. Snapshots of each trinucleotide-repeat system are provided in Fig. 2(a). Notably, systems rich in CG ‘stickers’ qualitatively show larger clusters than those lacking CG pairs. For example, in CGU, CAG, CUG, CCG, and CGG repeat systems a large fraction of the chains assemble to form distinctive clusters; with CAG clusters exhibiting roughly spherical shapes and CCG and CGG systems displaying more aspherical clusters. To better quantify the clustering behavior in the systems, we plot the fraction of chains in the largest cluster 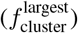 versus the cluster concentration, Fig. 2(b). Specifically, for CG-rich repeats, the vast majority of chains are concentrated within a single cluster. Additionally, there exists a strong positive correlation between the proportion of chains within the largest cluster and the concentration of clusters. This relationship suggests not only chain enrichment within CG-rich clusters but also dense packing of the chains within them. Thus, we hypothesize that CG-rich clusters should, therefore, exhibit slower dynamics. To verify this, we measure the mean square displacement (MSD) of individual chains in each of the systems, Fig. 2(c). In general, trinucleotide repeat systems that are deficient in CG stickers show roughly linear dependence of single-chain MSD with time (i.e., MSD *∝τ* ^1.0^). In contrast, as the CG content of the chains increases, we observe subdiffusive behaviors, where the MSD exponent decreases to 0.9, Fig. 2(c).

To further elucidate the origin of differences in clustering behaviors for the trinucleotide repeat sequences, we analyze the base-pairing propensities. Here, base-pairing propensities are primarily governed by two factors: (1) the strength of base pairs that each sequence supports [Fig. 3a] and (2) the relative number of repeat alignments that yield canonical/wobble base pairs [henceforth, referred to as base-pairing modes; insets in Fig. 3(b),(c)]. The former is enthalpic in nature, while the latter accounts for entropic contributions to binding.

**FIG. 3.**
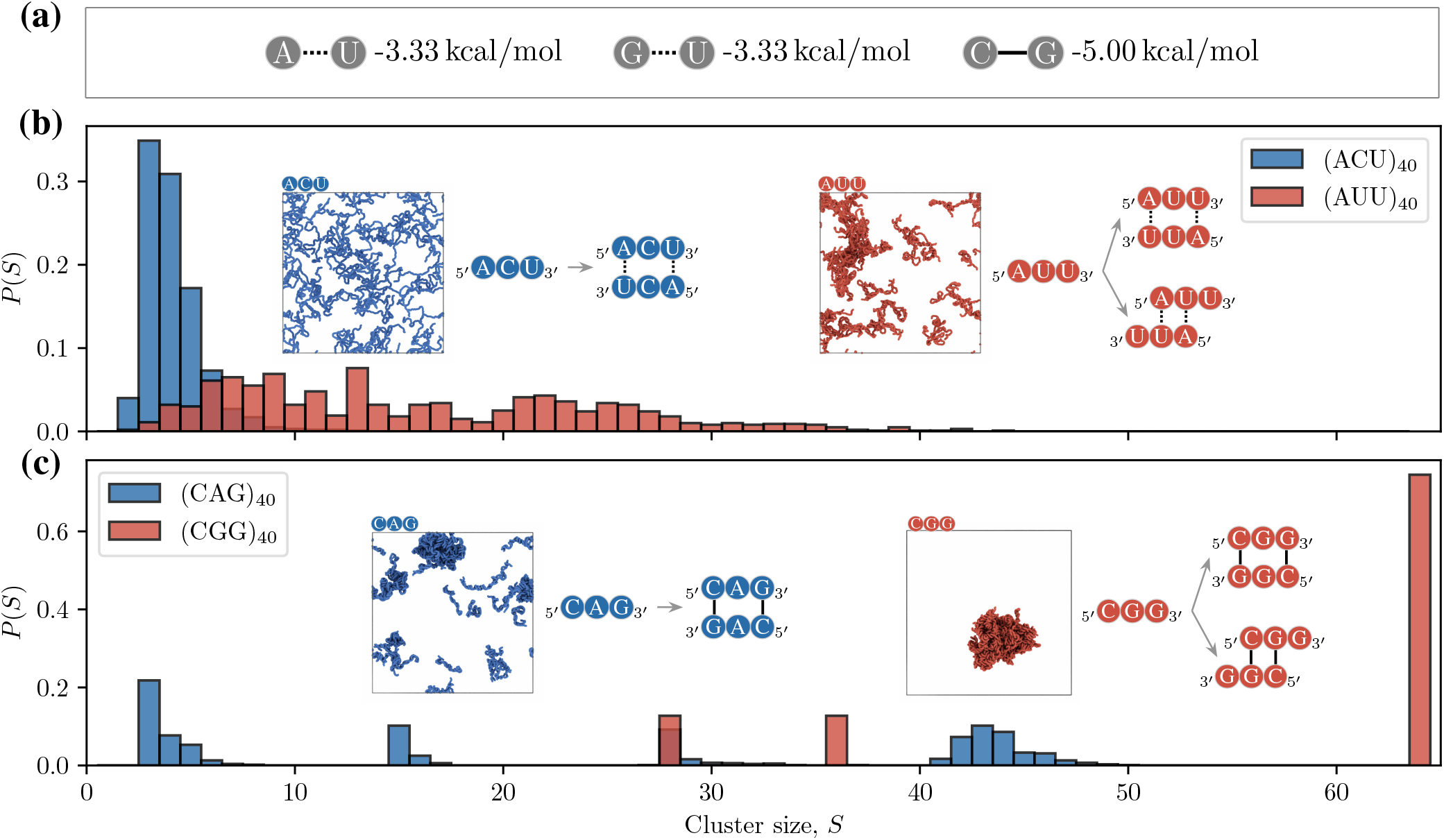
Entropic effects, via base-paring modes, contribute to stability of trinucleotide clusters. All systems are prepared at 200 *µ*M with 64 chains evenly distributed and melted at 373 K. Then the systems are cooled down to 293 K within 40 ns. Finally, the systems are simulated at 293 K for 2 *µ*s. The trajectories from the last 500 ns are used to measure the cluster size distributions.**(a)** Canonical and wobble base-pairing interaction strengths, as implemented in the coarse-grained model. **(b)** Comparison of cluster sizes between (ACU)_40_ and (AUU)_40_. Each repeat is able to form weaker A–U base pairs, but (AUU)_40_ has more probabilities to form base pairs. The simulation snapshots corresponding to the final state of each system, as well as the pairing modes for each system, are shown as insets. **(c)** Comparison of cluster sizes between (CAG)_40_ and (CGG)_40_. Each repeat is able to form stronger C–G base pairs, but (CGG)_40_ has more probabilities to form base pairs. The simulation snapshots corresponding to the final state of each system, as well as the pairing modes for each system, are shown as insets.

From an enthalpic standpoint, within the model, canonical (C–G, A–U) and wobble (G–U) base pairs are favoured, Fig. 3(a). Hence, in the’sticker–spacer’ framework [39, 40], C–G, A–U and G–U pairs can be regarded as stickers – complementary nucleotides that make sizable contributions to the stability of clusters. On the other hand, nucleotides that make negligible contributions to cluster stability can be regarded as spacers. By these definitions, (CAG)_47_ consists C–G stickers and A-spacer nucleotides. Broadly, the relative strengths of canonical and wobble base-pairing interactions (‘sticker– sticker’ binding) can explain the clustering propensities of trinucleotide sequences. For example, we compare the distribution of cluster sizes for select sequences in Fig. 3(b),(c). For repeats such as ACU, where C acts as a ‘spacer’ nucleotide, binding negligibly to either A or U, the cluster size distribution is governed by the interaction strength of the constituent A–U base pair. As a result, for repeats encoding weak base pair interactions (e.g., A–U in ACU and AUC) and distinct spacer nucleotides, the systems comprise many small clusters [Fig. 3(b)]. In contrast, similar repeats that instead encode stronger base-pair interactions (e.g., C–G in CAG and GAC) support larger clusters [Fig. 3(c)]. This dependency of clustering behaviors on base-pair energy is also supported by a recently proposed analytical sticker–spacer model for RNA repeats [31].

Interestingly, we also find a strong entropic contribution to the stability of trinucleotide clusters. Case in point, while both ACU and AUU encode identical sticker–sticker binding strengths, AUU on average supports larger clusters [Fig. 3(b)]. We find a similar behavior for CAG versus CGG for instance, where the latter on averages supports larger clusters [Fig. 3(c)]. This behavior naturally emerges in the simulations and can be explained on the basis of the entropic contributions to base pairing. Concretely, whereas ACU and CAG encode one dominant base-pairing mode, AUU and CGG encode two ideal base-pairing modes [Fig. 3(b),(c)]. The additional basepairing mode arises from the fact that both AUU and CGG lack a nucleotide that distinctively acts as a spacer and therefore multiple sequence alignments lead to favorable binding. Consequently, AUU and CGG clusters are on average larger than ACU and CAG clusters, respectively. Hence, repeats with multiple base-pairing modes provide additional entropic contributions to cluster formation – effectively reducing the free energy of clustering.

## Clustering propensities of trinucleotide repeats show qualitative agreement with experiments

One central focus of our work is probing whether basepairing tendencies are sufficient to explain differences in clustering behaviors of trinucleotide repeat sequences. In the previous section, we demonstrated that base-pairing strengths and the number of ideal base-pairing modes can collectively explain the trends observed in our simulations. A pertinent question arises: How accurately do the simulated trends reflect RNA clustering propensities observed in experiments? To this end, we turn to the work of Isiktas et al. [30], where they recently probed the ability of various trinucleotide repeat sequences to form foci (condensates) *in vivo*. Concretely, the authors expressed a given trinucleotide repeat in live cells and used green fluorescent protein (GFP) as probes to track the extent to which a given repeat sequence localized to condensates. Based on these experiments, they assessed and ranked the various trinucleotides according to their capacities to maintain condensates within live cells, identifying trinucleotides that resulted in the presence of more foci-positive cells as more conducive to condensation.

A key challenge in comparing such *in vivo* experiments with our simulations is the fact that, unlike our protein-free simulations, in addition to RNA, live cells also contain proteins which ultimately tune their ability to sustain condensates. Thus, differences in the number of condensate-positive cells could be attributed to variations in RNA–protein binding for a given trinucleotide repeat sequence, and not necessarily differences in RNA–RNA interactions. An advantage of simulations is that we can explicitly test the hypothesis: Are differences in base-pairing propensities sufficient to explain the variation in clustering for trinucleotide repeats? We, therefore, directly compare our simulation predictions with the trends captured in live cells. Specifically, we compare the fraction of chains in the largest cluster (simulations) with the fraction of foci-positive cells (experiments) [30], where both measurements are related to the propensities for RNA cluster formation, Fig. 4(a).

**FIG. 4.**
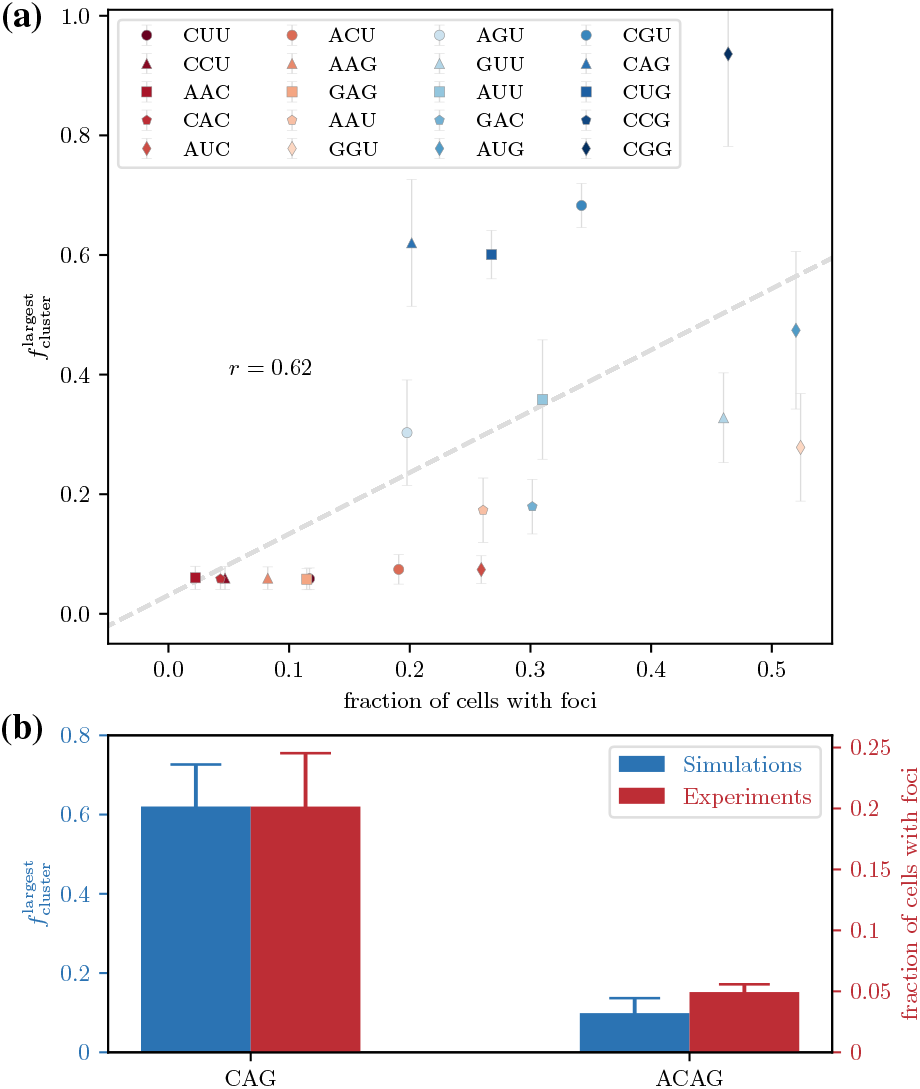
Comparison of RNA clustering in simulations with *in vivo* measurements of RNA-based condensates [30]. **(a)** Scatter plot of the fraction of foci-positive cells (experiments) and the fraction of the largest cluster 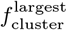 in our simulations. In our simulations, each RNA chain is composed of 40 nucleotide repeats while the *in vivo* experiment measured the localization (RNA foci) of RNA chains with each chain composing of 200 trinculeotide repeats. The error bars represent the standard deviation of the largest cluster fractions in our simulations. The dashed line represents linear fit of data and the Pearson coefficient is included. **(b)** Comparison of the fraction of the largest cluster 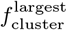 with the fraction of foci-positive cells for CAG and ACAG repeats. In our simulations, we simulate repeat sequences of (CAG)_40_ and (ACAG)_30_, where each sequence has the same length of 120 nucleotides. In the experiment, the localization (RNA foci) of (CAG)_200_ and (ACAG)_150_ were measured, where each sequence is has the same length of 600 nucleotides.

Here, we find a positive correlation between the fraction of RNA chains in the largest cluster in simulations and the fraction of foci-positive cells in the *in vivo* experiments [30] (Pearson correlation coefficient of 0.62), Fig. 4(a). Despite the reasonable agreement with experiments, there are, however, some disparities between the simulation predictions and experimental results. A notable exception occurs for (ACU)_40_ and (AUC)_40_, where the degree of clustering in the simulations is very low, despite the presence of foci in *in vivo* experiments for (ACU)_200_ and (AUC)_200_. To address potential finite size effects, we simulate (ACU)_200_ under the same conditions (Supplementary Fig. S7), which produces comparable results to the smaller system. Our analysis suggests that the disparities could be attributed to various other factors. Technically, this result may indicate a need for further calibration of the A–U interaction strength in the model, as well as the comparable G–U wobble base pair. Another plausible explanation for the discrepancy is the involvement of proteins inside cells, which often facilitate the formation of RNA foci, anticipated in the *in vivo* experiments. Notably, we expect trinucleotide repeats to nucleate proteins differently depending on their sequence and structure.

## Trinucleotide repeats show greater clustering propensities than analogue quadranucleotide repeat sequences

In addition to probing the clustering behavior of trinucleotide repeat sequences, we also assess the clustering propensities of quadranucleotide repeats. In particular, experimental work has demonstrated that odd (here, trinucleotide) and even (here, quadranucleotide) repeat sequences exhibit markedly different propensities for condensate formation [30], Fig. 4(b). Specifically, based on the *in vivo* experiments of (CAG)_200_ versus (ACAG)_120_ [30], we compare the clustering propensities of these systems in our simulations. For this comparison, we utilize (CAG)_40_ and (ACAG)_30_, which are comparable to the experimental systems, featuring sequences of the same length with odd and even repeats, respectively.

We find that whereas (CAG)_40_ chains stabilize large clusters [Fig. 5(a)], significantly smaller clusters are observed in the (ACAG)_30_ system [Fig. 5(a)]. This result suggest a greater clustering propensity for the odd-repeat (CAG) system compared to the even-repeat analogue (ACAG). We substantiate the generality of the observed differences in clustering tendencies between odd and even repeat sequences by also examining CUG versus UCUG [Fig. 5(b)]. Here, we observe a parallel behavior to that observed for CAG versus ACAG [Fig. 5(b)].

**FIG. 5.**
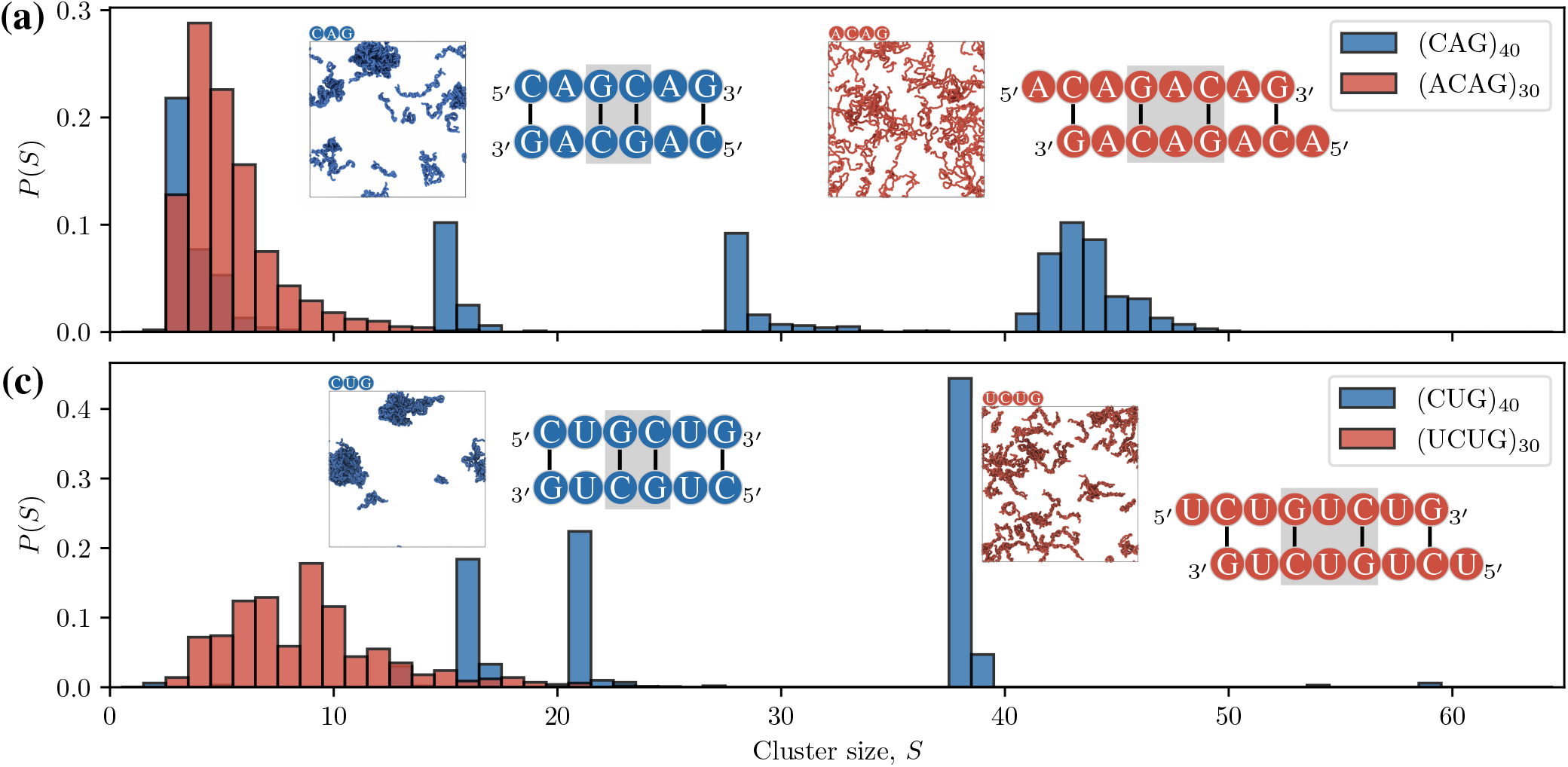
Comparison of clustering tendencies for trinucleotide (odd) and quadranucleotide (even) repeats. All systems are prepared at 200 *µ*M with 64 chains evenly distributed and melted at 373 K. Then the systems are cooled down to 293 K within 40 ns. Finally, the systems are simulated at 293 K for 2 *µ*s. The trajectories from the last 500 ns are used to measure the cluster size distributions. The simulation snapshots corresponding to the final state of each system, as well as the pairing modes for each system, are shown as insets. The main differences in pairing modes are highlighted by gray boxes. **(a)** The clustering propensities of (CAG)_40_ versus (ACAG)_30_. While (CAG)_40_ is able to form two consecutive C–G pairs that are separated by a ‘spacer’ A, the extra ‘spacer’ A in (ACAG)_30_ prevents it. **(b)** The clustering propensities of (CUG)_40_ versus (UCUG)_30_. Even though U is able to pair with G, the ideal pairing modes are dominated by strong C–G pairs. (CUG)_40_ is able to form two consecutive C–G pairs that are separated by ‘U’, the addition of extra ‘U’ in (UCUG)_30_ prevents it.

To assess the robustness of our predictions, we compare the fraction of chains in the largest cluster from our simulations against the fraction of foci-positive cells in the *in vivo* experiments for odd versus even repeats [Fig. 4(b)]. Here we find a good agreement between the simulation predictions and experiments, indicating that for odd versus even repeats, the model captures the differences in clustering behavior well.

Analyzing the ideal pairing modes for odd versus even repeat sequences provides insight into differences in clustering behaviors [inset in Fig. 5(a)]. The ratio of C–G base pairs to spacer (A) is 2:1 for CAG repeats, featuring two consecutive C–G pairs [grey shaded region, Fig. 5(a)]. In contrast, this ratio is 1:1 for ACAG repeats, which lack consecutive C–G pairs [inset, Fig. 5(a)]. Combining our simulation results with the experimental findings and predictions from an analytical model [30, 31] support the hypothesis that consecutive groups of at least two base pairs are required for homotypic RNA condensation.

## Balance between homotypic and heterotypic interactions govern cluster composition in binary mixtures

Finally, we also investigate binary mixtures composed of two different trinucleotide repeats to understand how competing interactions influence clustering propensities and the material properties of resulting clusters. In particular, we probe the clustering behaviors of three equimolar mixtures: (AAC)_40_–(AUG)_40_, (AAC)_40_–(CUG)_40_, and (CAG)_40_ (CUG)_40_, Fig. 6(a)–(c).

Concretely, in the single-component (AAC)_40_ system the largest cluster comprise about 4 chains – the system is essentially well-mixed [Fig. 6(a)]. In contrast, approximately 50% of the chains (ca. 32) are localized to the largest cluster in the single-component (AUG)_40_ system [Fig. 6(a)]. Mixing these two sequences results in 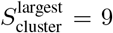 chains [Fig. 6(a)]. Interestingly, a significant fraction of the mixture comprises (AAC)_40_–(AUG)_40_ dimers. Since, AAC chains are more or less tethered to AUG, the mean square displacements (MSDs) of AAC and AUG in the mixture are identical.

**FIG. 6.**
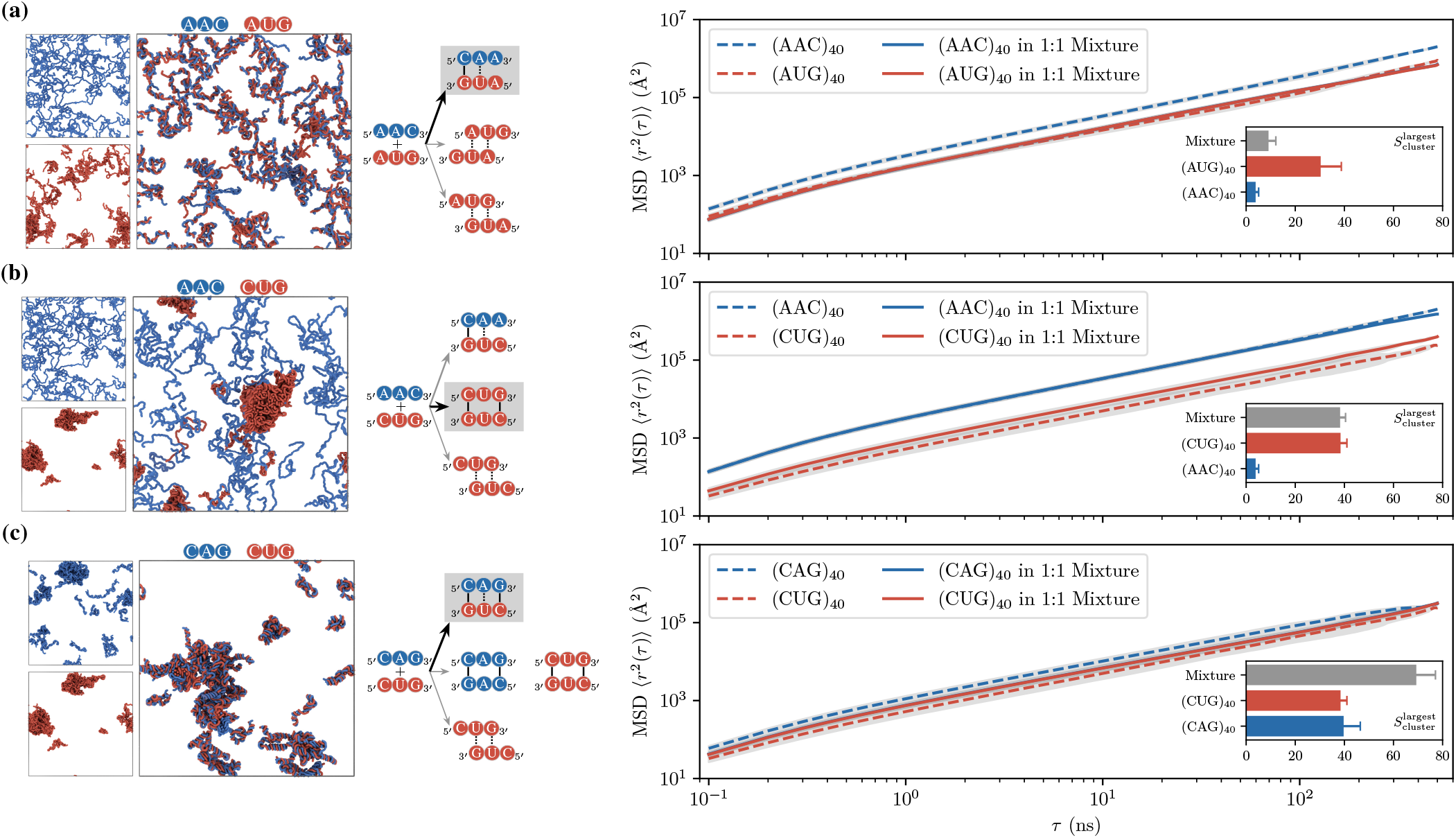
Material properties of RNA clusters in binary mixtures of trinucleotide repeats. Each equimolar mixture is composed of 64 + 64 = 128 chains and simulated with periodic boundary conditions at 200 *µ*M. The chains are evenly distributed and melted at 373 K. Then the systems are cooled down to 293 K within 40 ns. Finally, the systems are simulated at 293 K for 2 *µ*s. For comparison, the single component sysytems are simulated under the same condition with 64 chains. The trajectories from the last 500 ns are used to measure the cluster size distributions. For each subfigure, the snapshot of single-component systems and the mixtures, taken from the end of simulations, are shown on left with consistent coloring of different sequences. The pairing modes are shown in the middle with the minimum energy of pairing indicated by a gray box. The center of mass MSD (mean square displacement) of components in single-component systems and the mixture are show on right. In addition, the largest cluster sizes from simulations are shown as insets in the MSD subfigures. **(a)** Results for the binary mixture of (AAC)_40_ and (AUG)_40_. The dimers of (AAC)_40_ and (AUG)_40_ are dominant. **(b)** Results for the binary mixture of (AAC)_40_ and (CUG)_40_. Despite the ability of (AAC)_40_ and (CUG)_40_ to bind each other, (CUG)_40_ self-interactions mainly contribute to clustering, and (AAC)_40_ chains are almost excluded. **(c)** Results for the binary mixture of (CAG)_40_ and (CUG)_40_. With strong energetic compatibility, (CAG)_40_ and (CUG)_40_ are both contributing to clustering and experience relatively slow dynamics compared to other studied mixtures.

Analyzing the base-pairing strengths and modes reveal strong competition between heterotypic and homotypic interactions in regulating the clustering behavior of mixtures [middle panel, Fig. 6]. For a mixture of (AAC)_40_ and (AUG)_40_, the mixed pairing strength is stronger than the pairing of (AUG)_40_ by itself [middle panel, Fig. 6(a)]. On the other hand, self-pairing of (AUG)_40_ is entropically favored. The net effect is that the mixed pairing strength suppresses clustering relative to the single-component (AUG)_40_ system. Ultimately, the balance between the energetic and entropic terms, within a base-pairing framework, yields mainly dimers in this mixture.

The previous result is in stark contrast to that obtained for the (AAC)_40_–(CUG)_40_ mixture [Fig. 6(b)]. Here, (CUG)_40_ self-complementary pairing is dominant on both energetic and entropic grounds. Consequently, this mixture produces (CUG)_40_ clusters that almost entirely exclude (AAC)_40_ chains [Fig. 6(b)]. This behavior is also reflected in the dynamics of the chains in the mixture. Unlike the chains in the (AAC)_40_–(AUG)_40_ mixture [Fig. 6(a)], we find a clear separation of timescales for (AAC)_40_ and (CUG)_40_ in the (AAC)_40_– (CUG)_40_ mixture [Fig. 6(b)].

The findings in Fig. 6(a),(b) lead us to hypothesize that, in general, the compatibility of base-pairing strengths among trinucleotide repeats dictates the composition of RNA clusters in heterotypic systems. Here, compatibility refers to the similarity between the mixed pairing interactions and each of the self-complementary interactions. Notably, we find strong compatibility between such interactions in the (CUG)_40_–(CAG)_40_ mixture [Fig. 6(c)]. Additionally, the mixed pairing interaction is slightly stronger than each of the self-complementary ones. As a result, not only do the chains colocalize in clusters, but the overall clustering propensity of the system grows [Fig. 6(c)]. The largest cluster almost doubles in size. Moreover, the chains experience similar dynamics in the mixture relative to the single-component (CUG)_40_ and (CAG)_40_ systems.

## DISCUSSION

We have successfully implemented a coarse-grained model [15] to investigate the clustering behaviors of RNA sequences. In this model, each RNA nucleotide is represented by a unique bead, and nucleotide–nucleotide interactions are adjusted to capture the relative strengths of canonical (C–G, A–U) and wobble (G–U) base pairs. Additionally, the model incorporates a many-body base-pairing potential previously optimized to capture the A-form helical structure of CAG-repeat sequences and reproduce the length-dependent clustering behaviors observed in such sequences. However, employing many-body potentials can lead to computational inefficiencies, even with nucleotide-level coarse-graining. To enhance efficiency and broaden the accessibility of this method, we explicitly derive the forces for the many-body potential (66 force components in total) and implement them in the LAMMPS software [26]. This implementation allows us to leverage the excellent parallelization capabilities of LAMMPS.

In a series of tests, we demonstrate the correctness of our implementation. These tests include comparing the numerical integrals to the analytical ones via the the one-side Euler method, assessing energy conservation in the microcanonical ensemble, as well as reproducing single and multi-chain properties of (CAG)_*n*_ repeat systems [15]. In addition to ensuring correctness and stability, our implementation notably enhances simulation efficiency, achieving speedups of 3 to 5 times faster in our test cases. Furthermore, we demonstrate the scalability of our LAMMPS implementation; increasing the number of computing cores results in near-linear scaling.

Leveraging the increased efficiency of the model, we systematically explore the clustering propensity of all 20 non-redundant trinucleotide repeats and compare the results with experimental data. Overall, we observe a positive correlation between simulations and *in vivo* experiments [30]. Interestingly, our simulations reveal significant entropic contributions to the stabilization of RNA clusters, particularly considering the degeneracy of ideal base-pairing modes each repeat encodes. Remarkably, these effects emerge organically in the simulations.

Expanding our investigation to various RNA sequences, we compare the clustering behavior of odd versus even repeats, focusing on CAG versus ACAG and CUG versus UCUG repeats. While the CAG and CUG repeats show strong clustering, the addition of an extra ‘A’ or ‘U’ spacer for the ACAG and UCUG repeats, respectively, suppresses cluster formation. These results are also consistent with experiments [30]. Furthermore, these findings underscore the importance of consecutive groups of at least two base pairs for homotypic RNA phase separation [31].

Additionally, our examination of binary mixtures of trinucleotide repeats reveals that RNA cluster formation heavily depends on the balance between mixed base-pairing interactions (heterotypic) and self-complementary interactions (homotypic). We observe that this equilibrium governs both the size and composition of clusters. In scenarios where heterotypic and homotypic interactions strongly compete, dimers may form (as in the (AAC)_40_–(AUG)_40_ mixture). Conversely, when the self-complementary interactions of one repeat sequence predominate, homotypic RNA clusters emerge (as observed in the (AAC)_40_–(CUG)_40_ mixture). Finally, in cases where both types of interactions significantly contribute and are compatible energetically and entropically, such systems exhibit co-clustering of RNA trinucleotide repeats (as in the (CAG)_40_–(CUG)_40_ mixture).

We expect that the observed trends in clustering behaviors reported here may also apply to similar biopolymers. However, the studied sequences represent only a small fraction of the vast sequence space. Typically, mutations at a few sites in such biopolymers can significantly alter their clustering behaviors. Hence, our implementation of the coarse-grained RNA model opens avenues for the exploration of a more diverse range of sequences.

It is essential to emphasize that in the current implementation solely considers Watson–Crick canonical base pairs and G–U wobble base pairs, as they are the most prevalent according to an analysis of experimental RNA structures (Supplementary Fig. S1, Supplementary Table S2). Our findings underscore that while accounting for canonical and wobble base-pairing propensities enables qualitative predictions of clustering tendencies in trinucleotide RNA repeats, for precise quantitative predictions, other interactions such as noncanonical base pairs and base stacking may be indispensable.

Additionally, in the current model, interaction strengths represent effective free energies assuming charge neutralization. Given that ions such as Mg^2+^ play crucial roles in RNA condensation, explicitly considering ion effects (and electrostatic effects in general) [41, 42] should enhance the accuracy of predictions regarding trinucleotide clustering. For instance, G-rich sequences also yield stable RNA clusters, likely sustained by a synergy between non-canonical base-pairing and ionic effects [43, 44]. Moreover, meticulous incorporation of electrostatic effects paves the way for quantitative analysis of interactions between such RNA sequences and proteins [45–51]. Lastly, expanding these models to encompass a wider array of RNA secondary structures could enhance their utility – both in protein-free solutions and protein–RNA mixtures, where structural recognition could be important.

Overall, the evolving approach in developing efficient RNA phase separation models, which explicitly account for sequence-dependent effects will enhance our ability to correlate RNA sequences with macroscopic condensate features. When integrated with experimental data, these biophysical models contribute to advancing our understanding of spatiotemporal organization within cells, shedding light on mechanisms involved in abnormal condensation, and elucidating principles for designing soft materials based on RNA compartmentalization.

## METHODS

### A. The coarse-grained model of RNA

In the single-interaction-site (SIS) model [15], each RNA nucleotide is represented by a single bead. As the center of coarse-graining is not specified in the original implementation, we utilize the C3’ atom on ribose to determine the location of the coarse-grained bead. This choice is deemed reasonable based on the RMSD comparison against the CAG repeat crystal structure, Supplementary Fig. S2, and our analysis of the force field parameters against bioinformatics data [52], Supplementary Fig. S1.

In this model, the total potential energy (*U*) is decoupled into three coarse-grained effective potentials:

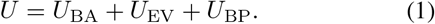

A schematic representation of the potential terms is presented in Fig. 1(a). Here, *U*_BA_ describes the bonded interactions of RNA nucleotides, consisting of bond and angle restraints between the connected beads. The specific form of the potential is:

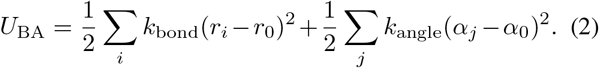

The excluded volume interaction *U*_EV_ is given by the Weeks– Chandler–Andersen potential:

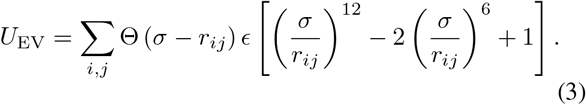

Θ is the Heavyside step function and if the two beads are on the same chain and involved in bond or angle potentials, then this term is not computed. For the short-ranged interactions that mimic the canonical Watson–Crick A-form base-pairing between beads *i* and *j*, the base-pair potential, *U*_BP_ is given as:

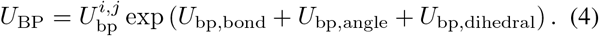

Here 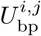 is the interaction strength between two base-pairs that depends on nucleotide types of bead *i* and *j*. In the current model, only canonical A–U, C–G and G–U wobble pairs are considered. The specific forms for each term are listed as follows:

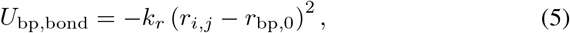

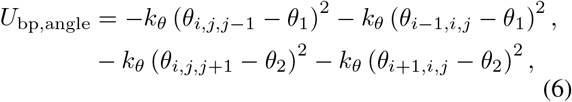

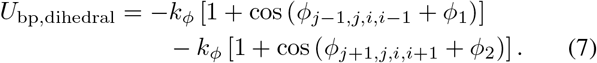

The interaction strength 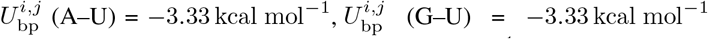, and 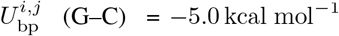. These interaction strengths were selected to correspond to the number of hydrogen bonds formed between the base pairs, comprising two H-bonds for A–U and G–U, and three for G–C.

We confirm that the other parameters of the base-pairing interaction potential align well with the A-form structure by analyzing approximately 500 RNA A-form antiparallel structures (Supplementary Table S2) from the Nucleic Acid Knowledgebase (NAKB) [52]. Our analysis reveals no significant differences in the distributions of bond distances, angles, and dihedrals, Supplementary Fig. S2. As a result, in our LAMMPS implementation, all other bonded parameters for the A–U, G– C, and G–U pairs remain consistent with the original model. All parameter values for the potentials discussed herein are detailed in Supplementary Table S1.

### B. Implementation of RNA model in LAMMPS Software

Implementation of custom potentials in LAMMPS requires explicit coding of the corresponding forces. Therefore, we first derive the force terms of the base-pair potential in the model (Eq. 4, 5, 6, 7). The force calculation for base pairing requires the careful evaluation of forces due to multiple angle references. We use the formalism that is described in Ref. [27] to derive all 66 force components.

Below is one example of our derivation for calculating the 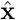 component of the force **f**_*i*_ on bead *i* at position **r**_*i*_ = (*x*_*i*_, *y*_*i*_, *z*_*i*_) due to the dihedral angle *ϕ*_*j*+1,*j*,*i*,*i*+1_ [Fig. 1(a), bottom panel]. First, the displacement vectors are defined from the position vectors of coarse-grained beads **r**_*i*+1_, **r**_*i*_, **r**_*i−*1_, **r**_*j*+1_, **r**_*j*_, and **r**_*j−*1:_

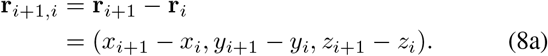

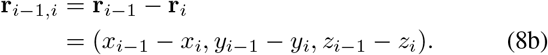

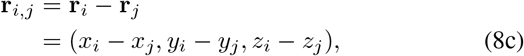

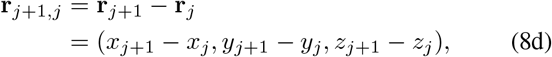

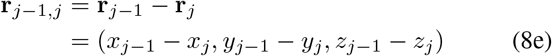

Then, the normal vectors of the surfaces that are formed by these vectors are defined as

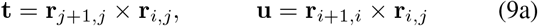

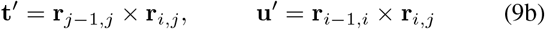

Thus, the angles can be calculated using the cos function

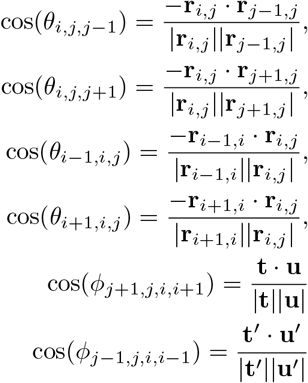

The distance and angles can be used to calculate the basepair potential *U*_BP_ (Eq.4) and thus the force can be derived:

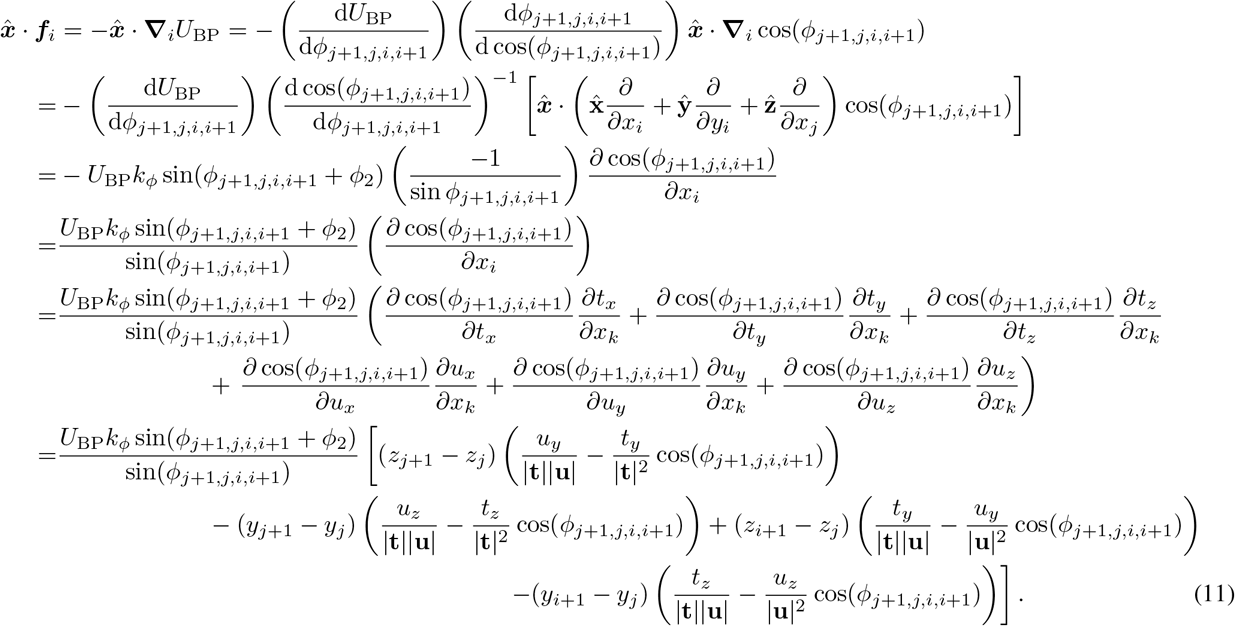

A similar derivation procedure applies to calculations of all other 65 force components. For the pressure calculation, LAMMPS uses the pressure tensor that takes into account the contribution of the kinetic energy tensor and the virial tensor.

Thus, for example, the contribution of the 4 forces **f**_*j*+1_, **f**_*j*_, **f**_*i*_ and **f**_*i*+1_ from the dihedral angle *ϕ*_*j*+1,*j*,*i*,*i*+1_ to the *xx* component of the virial tensor is

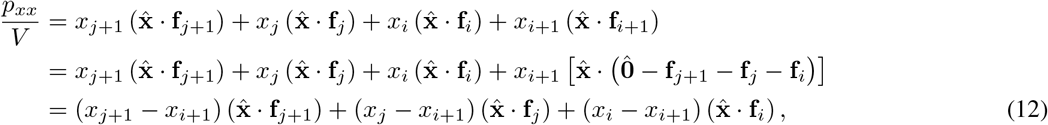

where, *V* is the total volume and 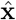 is the unit vector of the *x* direction in the Cartesian coordinate system. Similarly, the contributions to the pressure tensor are calculated for all the forces involved in the base-pair interaction.

Finally we implemented the forces and the tensors. By compiling with the force-field files (instructions are included in Github), the base pair potential can be accessed in LAMMPS by calling base/rna, with the freedom to set the interaction strength 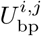, equilibrium distance *r*_0_, the equilibrium angles *θ*_*i*,*j*,*j−*1_, *θ*_*i−*1,*j*,*j*_, *θ*_*i*,*j*,*j*+1_, *θ*_*i*+1,*j*,*j*_, and the dihedrals *ϕ*_*j−*1,*j*,*i*,*i−*1_, *ϕ*_*j*+1,*j*,*i*,*i*+1_, as well as their coefficients *k*_*r*_, *k*_*θ*_ and *k*_*ϕ*_ to any values depending on the systems of interest (9 parameters). An example script for defining this RNA model in LAMMPS is included in the Supplementary material (Listing S1).

### Simulation

For the microcanonical simulation of the single chain (AGGCAGCAGAAAAGACGACCCA), a timestep of 10 fs is employed. The potential energy and the kinetic energy are computed every 1 ps. The average drift of the total energy |Δ*E*_total_| _*t*_ is calculated by dividing the absolute overall drift from the initial total energy |*E*_total_(*t* + Δ*t*) *− E*_total_(*t*)| by the duration of time Δ*t*:

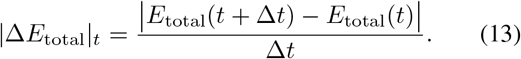

For all the canonical ensemble simulations of RNA repeats, the timestep is set the same as the simulations in the microcanonical ensemble (10 fs). In multichain simulations, each system is composed of 64 chains, except where otherwise specified, and simulated with periodic boundary conditions. The systems are prepared at 200 *µ*M concentration with chains evenly distributed in the initial configuration. For (AUC)_200_ we prepared the system with 27 chains at the same number density.

For all *NV T* simulations, a Langevin thermostat with low friction is used (damping time = 100,000 timesteps, with the integration step size of 10 fs). First, the chains are melted at 373 K for 5 ns. This ensures that most of the chains are in an unfolded state. Then the temperature is reduced by 10 K, and the system is simulated with the Langevin thermostat for 5 ns. Repeating this cooling procedure, the system is annealed to 293 K within 40 ns. Finally, the systems are simulated at 293 K for 2 *µ*s. Configurations for trajectory files are recorded every 0.1 ns. For estimating the average quantities near equilibrium, the data from the last 500 ns are used.

### Testing the scalability of the model

For the benchmarking results presented in Fig. 1(d,e), we prepare a compressed condensate of (CAG)_47_ with 64 chains and simulate the system for 100,000 timesteps with a Langevin thermostat at 293 K. A timestep of 10 fs is used for our GPU and CPU tests. Then we obtain the simulation speed by dividing the total number of steps by wall clock time.

For further testing, we use the original OpenMM code and simulate the system using NVIDIA A100 Tensor Core 80GB graphics cards on Della cluster (Princeton Research Computing). We use OpenMM version 7.5 for code compatibility and vary the number of GPUs and the precision to analyze the scalability.

Finally, to test the scalability of our implementation, we simulate the same system in LAMMPS using Intel Cascade Lake CPUs on Della cluster (Princeton Research Computing), and vary the number of CPUs from 1 to 128.

### Cluster measurements

For defining clusters, we used the criteria based on the spatial distances of individual chains. Specifically, two chains are in the same cluster if two of their nucleotides are within 1.2 *r*_0_, where *r*_0_ = 13.8 Å is equilibrium base-pairing distance. We obtained the cluster data with gyration tensors using Ovito [53] version 3.10. For the concentrations of clusters, the gyration tensor is used, which is diagnonalized for a cluster as

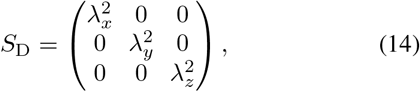

where the principle moments 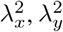 and 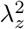 describes effective extension of the cluster in the principal axial directions. Thus, the volume of the cluster can be estimated using an ellipsoid as 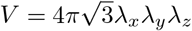.

## ACKNOWLEDGEMENTS

The simulations reported on in this manuscript was substantially performed using the Princeton Research Computing resources at Princeton University, which is a consortium of groups led by the Princeton Institute for Computational Science and Engineering (PICSciE) and Office of Information Technology. We thank Nathaniel Hess for helpful discussions regarding the implementation of custom potentials in LAMMPS and the manuscript. We also thank Dr. Alina Emelianova, Ananya Chakravarti and Prof. Athanassios Panagiotopoulos for helpful feedback on the manuscript. This research was supported by departmental start-up funds allocated to J.A.J. through the Department of Chemical and Biological Engineering and the Omenn-Darling Bioengineering Institute at Princeton University. This research was also partially supported by the National Science Foundation (NSF) through the Princeton University (PCCM) Materials Research Science and Engineering Center DMR-2011750. The authors also acknowledge funding from the Silicon Valley Community Foundation.

## CODE AND DATA AVAILABILITY

The source code for running the RNA model in LAMMPS, input scripts, as well as instructions for compiling can be found at https://doi.org/10.5281/zenodo.10428292 and at Joseph Group GitHub repository: https://github.com/josephresearch/RNA_lammps.

## Notes

https://github.com/josephresearch/RNA_lammps

### Competing Interest Statement

The authors have declared no competing interest.

### Summary of Updates

More benchmark performed; text and figure contents updated.

