## Supplementary file for "Accelerated simulations of RNA clustering: a systematic study of repeat sequences"

### Supporting Information for: Accelerated simulations of RNA clustering: a systematic study of repeat sequences

#### A. Parameters

TABLE S1. Parameters for the coarse-grained RNA model

| Parameter | Value | Parameter | Value | Parameter | Value |
|---|---|---|---|---|---|
| $k_{\rm bond}$ | $15.0\,{\rm kcal\ mol^{-1}Å^{-2}}$ | $r_0$ | $5.9\,{\rm Å}^2$ | $k_{\rm angle}$ | $10.0\,{\rm kcal\ mol^{-1}rad^{-2}}$ |
| $\alpha_0$ | $2.618\,{\rm rad}$ | $\sigma$ | $10\,{\rm Å}$ | $\epsilon$ | $2.0\ {\rm kcal\ mol^{-1}}$ |
| $U_{\rm bp}^{i,j}$ (C-G) | $-5.00\,{\rm kcal\ mol^{-1}}$ | $U_{\rm bp}^{i,j}$ (A-U) | $-3.33\,{\rm kcal\ mol^{-1}}$ | $U_{\rm bp}^{i,j}$ (G-U) | $-3.33\,{\rm kcal\ mol^{-1}}$ |
| $k_r$ | $3.0\,{\rm kcal\ mol^{-1}Å^{-2}}$ | $r_{\rm bp,0}$ | $13.8\,{\rm Å}^2$ | $k_\theta$ | $1.5\,{\rm kcal\ mol^{-1}rad^{-2}}$ |
| $\theta_1$ | $1.8326\,{\rm rad}$ | $\theta_2$ | $0.9425\,{\rm rad}$ | $k_\phi$ | $0.5$ |
| $\phi_1$ | $1.8326\,{\rm rad}$ | $\phi_2$ | $1.1346\,{\rm rad}$ | | |

##### B. Bioinformatic Survey of RNA Structures

We first coarse-grain atomistic structures (C3' as heavy atom center) to calculate base-pair distances and the relative angles. As shown in Fig. S1(a), the distribution of base-pairing distance for all three A–U, C–G, and G–U pairs are indeed close to the reference distance $r_0 = 13.8\,\text{Å}$ in the model. For the bond and dihedral angles formed by these pairs, we also combine these terms into three groups and the plot corresponding scatter plots (Fig. S1 (b), (c), (d)). In addition, we also define the corresponding energy functional forms of these three groups as follows:

$$f_1(\theta_{i,j,j-1}, \theta_{i-1,j,j}) = \exp\left[-k_\theta(\theta_{i,j,j-1} - \theta_1)^2 - k_\theta(\theta_{i-1,i,j} - \theta_1)^2\right], \tag{S1}$$

$$f_2(\theta_{i,j,j+1}, \theta_{i+1,j,j}) = \exp\left[-k_\theta(\theta_{i,j,j+1} - \theta_2)^2 - k_\theta(\theta_{i+1,i,j} - \theta_2)^2\right], \tag{S2}$$

$$f_3(\phi_{j-1,j,i,i-1}, \phi_{j+1,j,i,i+1}) = \exp\left[-k_\phi(1 + \cos(\phi_{j-1,j,i,i-1} + \phi_1)) - k_\phi(1 + \cos(\phi_{j+1,j,i,i+1} + \phi_2))\right]. \tag{S3}$$

Thus, the base-pair energy can be expressed as $U_{\rm BP} = U_{\rm bp}^{i,j} \exp(U_{\rm bp,bond}) f_1 f_2 f_3$. As shown in Fig. S1, the distributions of experimental values align well with the corresponding functional forms computed using the same references.

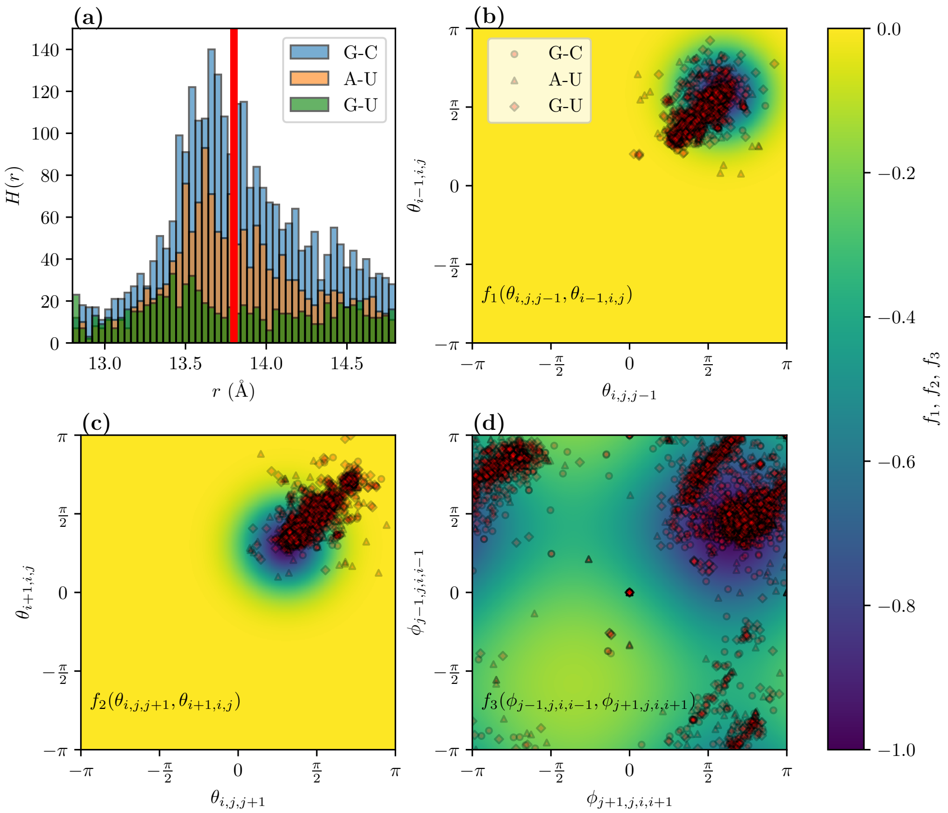


FIG. S1. **Evaluating current parameters of base-pairing potential against experimental structures.** **(a)** The base-pair distance (C3' ⋯ C3') distributions for the G–C, A–U, and G–U base-pairs peak close to the equilibrium distance in the model (red line, $r_0 = 13.8\,\text{Å}$). **(b)** Heatmap of base-pair angles $\theta_{i,j,j-1}$ and $\theta_{i-1,j,j}$, and the related base-pair functional form $f_1(\theta_{i,j,j-1}, \theta_{i-1,j,j})$. The scatter plot of the experimental values coincides with the minima of the potential for all three base pairs. **(c)** Heatmap of base-pair angles $\theta_{i,j,j+1}$ and $\theta_{i+1,j,j}$, and the related base-pair functional form $f_2(\theta_{i,j,j+1}, \theta_{i+1,j,j})$. The bioinformatic data coincides well with the corresponding functional form. **(d)** Heatmap of the base-pair dihedral angles $f\phi_{j-1,j,i,i-1}$ and $\phi_{j+1,j,i,i+1}$, and the related base-pair functional form $f_3(\phi_{j-1,j,i,i-1}, \phi_{j+1,j,i,i+1})$. The overlay of bioinformatic data shows good agreement with the energy landscape of the functional form.

TABLE S2. **Surveyed PDB IDs of RNAs in A-form**. Data obtained from the Nucleic Acid Knowledgebase (NAKB) [1] with options of (1) antiparallel double helices, (2) A-form RNA and (3) without proteins.

| | | | | | | | | | | | | | | | |
|---|---|---|---|---|---|---|---|---|---|---|---|---|---|---|---|
| 1AJF | 1AL5 | 1ATO | 1ATV | 1ATW | 1BZ2 | 1BZT | 1BZU | 1CSL | 1D0U | 1D4R | 1DQF | 1DQH | 1DUQ | 1EBQ | 1EBR |
| 1EKA | 1EKD | 1ELH | 1ESH | 1F6X | 1F79 | 1F7F | 1F7I | 1FYP | 1G2J | 1G3A | 1GUC | 1HS1 | 1HS2 | 1HS3 | 1HS4 |
| 1HS8 | 1I3X | 1I3Y | 1I46 | 1I4B | 1I9X | 1IDV | 1JZC | 1JZV | 1K4B | 1K5I | 1KFO | 1L1W | 1L3Z | 1LC6 | 1LMV |
| 1LNT | 1MFK | 1MIS | 1MNX | 1MUV | 1MV1 | 1MV2 | 1MV6 | 1MWG | 1N8X | 1NA2 | 1NC0 | 1NLC | 1NZ1 | 1O3Z | 1OQ0 |
| 1OW9 | 1PBM | 1PJY | 1QBP | 1QC0 | 1QCU | 1QES | 1QET | 1RNA | 1RXA | 1RXB | 1SA9 | 1SAQ | 1SDR | 1SLO | 1SLP |
| 1SY4 | 1SYZ | 1SZY | 1T0D | 1T0E | 1TJZ | 1TUT | 1U2A | 1WTS | 1WTT | 1WVD | 1XHP | 1XV0 | 1Y6S | 1Y6T | 1Y73 |
| 1Y90 | 1Y95 | 1Y99 | 1YFV | 1YLG | 1YN2 | 1YNC | 1YY0 | 1YZD | 1Z30 | 1Z79 | 1Z7F | 1ZIH | 1ZX7 | 1ZZ5 | 205D |
| 255D | 259D | 280D | 283D | 2A04 | 2A0P | 2AHT | 2AO5 | 2B7G | 2B8R | 2CD1 | 2CD3 | 2CD5 | 2DD2 | 2ES5 | 2ET3 |
| 2FQN | 2G1G | 2G3S | 2G91 | 2G92 | 2GBH | 2GIP | 2GVO | 2H49 | 2HNS | 2JR4 | 2JRG | 2JSG | 2JUK | 2JWV | 2JXQ |
| 2JXS | 2K41 | 2K5Z | 2K65 | 2K66 | 2K7E | 2KD8 | 2KEZ | 2KF0 | 2KOC | 2KPC | 2KPD | 2KRQ | 2KRV | 2KRZ | 2KVN |
| 2KXZ | 2KY0 | 2KY1 | 2KYD | 2KYE | 2L2J | 2L6I | 2L8F | 2L8U | 2L8W | 2L9E | 2LAC | 2LBQ | 2LK3 | 2LP9 | 2LPA |
| 2LPS | 2LPT | 2LV0 | 2LWK | 2M1O | 2MER | 2MS5 | 2MVS | 2MVY | 2MXJ | 2MXK | 2N4L | 2NBY | 2NCR | 2O32 | 2O3W |
| 2O3Y | 2O81 | 2O83 | 2OE5 | 2OE6 | 2OE8 | 2Q1O | 2Q1R | 2QEK | 2R1S | 2R20 | 2R21 | 2R22 | 2RLU | 2RO2 | 2RPT |
| 2U2A | 2V6W | 2V7R | 2VAL | 2VUQ | 2W89 | 2XSL | 2Y95 | 333D | 353D | 397D | 3BNL | 3BNT | 3CGP | 3CGQ | 3CGR |
| 3CGS | 3CJZ | 3FTM | 3GLP | 3GVN | 3HGA | 3JXQ | 3JXR | 3LOA | 3MEI | 3ND4 | 3NJ6 | 3NJ7 | 3OK2 | 3P4B | 3P4C |
| 3P4D | 3R1C | 3R1D | 3R1E | 3S49 | 3SZX | 3TD0 | 402D | 405D | 409D | 413D | 420D | 433D | 434D | 435D | 438D |
| 439D | 462D | 464D | 466D | 472D | 4A4R | 4A4S | 4E48 | 4E59 | 4E5C | 4E6B | 4J50 | 4JAB | 4JRT | 4K27 | 4K31 |
| 4KYY | 4MCE | 4MCF | 4MS9 | 4MSB | 4MSR | 4NFO | 4NFP | 4NFQ | 4O41 | 4P3T | 4P3U | 4P43 | 4PCJ | 4RBQ | 4RBY |
| 4RBZ | 4RC0 | 4U34 | 4U35 | 4U37 | 4U38 | 4U3L | 4U3O | 4U3P | 4U3R | 4U47 | 4U78 | 4XW0 | 4YN6 | 4ZC7 | 5AY2 |
| 5AY3 | 5AY4 | 5BTM | 5C5W | 5D8T | 5DA6 | 5DER | 5DHB | 5DHC | 5DO5 | 5EW4 | 5EW7 | 5HBW | 5HBX | 5HBY | 5HN2 |
| 5HNJ | 5HNQ | 5KRG | 5KVJ | 5L00 | 5LQO | 5LQT | 5LR3 | 5LR4 | 5LR5 | 5LSN | 5MWI | 5NXT | 5T3K | 5TDJ | 5TGP |
| 5TKO | 5UED | 5UEE | 5UEF | 5UEG | 5UF3 | 5UZ6 | 5V0H | 5V0J | 5V0O | 5V1K | 5V1L | 5V2H | 5V2R | 5V9Z | 5VH7 |
| 5VH8 | 5VR4 | 5WQ1 | 5ZEM | 6AAS | 6AAU | 6AZ4 | 6BG9 | 6BGB | 6BMD | 6C8D | 6C8E | 6C8I | 6C8J | 6C8K | 6C8M |
| 6C8N | 6C8O | 6CAB | 6CXZ | 6CY2 | 6CY4 | 6E7L | 6EZ0 | 6H0R | 6HBX | 6HC5 | 6HMO | 6HU6 | 6I1V | 6I1W | 6IA2 |
| 6JBG | 6L0Y | 6MXQ | 6N8F | 6N8H | 6N8I | 6NOA | 6U6J | 6U79 | 6U89 | 6U8U | 6UGI | 6UGJ | 6VA1 | 6VA2 | 6VA3 |
| 6VA4 | 6VEM | 6VU1 | 6VZC | 6WY2 | 6WY3 | 6XUR | 6XUS | 6XWJ | 6XWW | 6XXB | 6YMC | 6Z18 | 6ZPF | 6ZQ9 | 6ZR1 |
| 6ZRS | 6ZX5 | 6ZX8 | 7A3Y | 7A9N | 7A9O | 7A9Q | 7A9S | 7A9T | 7ECJ | 7ECK | 7ECL | 7ECM | 7ECN | 7ECO | 7ECP |
| 7EDT | 7EDU | 7JU1 | 7KUB | 7KUC | 7KUD | 7KUK | 7KUM | 7KUN | 7KUO | 7KUP | 7LNE | 7LNF | 7LNG | 7QTN | 7QUA |
| 7U87 | 7U88 | 7U89 | 7U8A | 7U8B | 7UME | 7VFT | 7Y2B | 7Y2P | 7Y8P | 8AMG | 8AMI | 8AMJ | 8AMK | 8AML | 8AMM |
| 8AMN | 8BWT | 8CLM | 8CLR | 8FCS | 8I43 | 8SWG | 8SWO | 8SX5 | 8SX6 | 8SXL | 8SY1 | | | | |

##### C. Implementation of RNA model in LAMMPS

The following are the potential energy definitions in a LAMMPS input script.

```
units real
dimension 3
boundary p p p
atom_style full
#### harmonic bond between bonds
bond_style harmonic
#    coeff for harmonic bond, specify 3:
#    * bond type
#    * K/2
#    * equilibrium distance
bond_coeff 1 7.5 5.9

#### harmonic angle potential between bonds
angle_style harmonic
#    coeff for harmonic angle, specify 3:
#    * angle type
#    * K/2
#    * equilibrium angle in degrees
angle_coeff 1 5.0 150.0

#### overlay of WCA-LJ potential and Base-Pair Potential
pair_style hybrid/overlay lj/cut 10.0 base/rna 18.0

#  pair_coeff for WCA-LJ potential, specify 2:
#    * energy
#    * sigma
pair_coeff * * lj/cut 2.0 8.9089871814
pair_modify pair lj/cut shift yes

#  pair_coeff for Base Pair Interactions, specify 9:
#    * energy Ubp: A-1, C-2, G-3, U-4
#    * harmonic K_r
#    * Equilibrium distance r_0
#    * harmonic K_theta
#    * dihedral K_phi
#    * Equilibrium angle theta1
#    * Equilibrium angle theta2
#    * Equilibrium angle phi1
#    * Equilibrium angle phi2
pair_coeff 1 1 base/rna 0.00000 3.0 13.8 1.5 0.5 1.8326 0.9425 1.8326 1.1345 #AA
pair_coeff 1 2 base/rna 0.00000 3.0 13.8 1.5 0.5 1.8326 0.9425 1.8326 1.1345 #AC
pair_coeff 1 3 base/rna 0.00000 3.0 13.8 1.5 0.5 1.8326 0.9425 1.8326 1.1345 #AG
pair_coeff 1 4 base/rna 3.33333 3.0 13.8 1.5 0.5 1.8326 0.9425 1.8326 1.1345 #AU
pair_coeff 2 2 base/rna 0.00000 3.0 13.8 1.5 0.5 1.8326 0.9425 1.8326 1.1345 #CC
pair_coeff 2 3 base/rna 5.00000 3.0 13.8 1.5 0.5 1.8326 0.9425 1.8326 1.1345 #CG
pair_coeff 2 4 base/rna 0.00000 3.0 13.8 1.5 0.5 1.8326 0.9425 1.8326 1.1345 #CU
pair_coeff 3 3 base/rna 0.00000 3.0 13.8 1.5 0.5 1.8326 0.9425 1.8326 1.1345 #GG
pair_coeff 3 4 base/rna 3.33333 3.0 13.8 1.5 0.5 1.8326 0.9425 1.8326 1.1345 #GU
pair_coeff 4 4 base/rna 0.00000 3.0 13.8 1.5 0.5 1.8326 0.9425 1.8326 1.1345 #UU

special_bonds lj 0.0 0.0 1.0 # no excluded volume calculation for bonded beads
```

Listing S1. The part of the potential definitions in LAMMPS scripts.

##### D. Validation of results for CAG repeats

We first validate the single chain simulation results for a short $(CAG)_2$ duplex as shown in Figure. S2. For comparison with the crystal structure (PDB code: 3NJ6), the modified singe chain (AGGCAGCAGAAAAGACGACCCA) is simulated using the same Langevin thermostat at 293 K for 500 ns. The structure is recorded every 10 ps. The RMSD values are calculated against the C3' atoms in the 3NJ6 reference structure. Specifically, the calculation is performed using the Python API from PyMOL [2] version 2.5, and data from the last 400 ns is used to calculate the average.

In addition to the verification for single chains, we simulate multichain systems of RNA $(CAG)_n$ repeats to test their length-dependent phase behavior. Specifically, we simulate two CAG repeat systems, $(CAG)_{47}$ at concentration 200 $\mu$M, and $(CAG)_{20}$ at concentration 50 $\mu$M. Each system is composed of 64 chains and simulated with periodic boundary conditions. The systems are prepared with an even distribution of chains and melted at 373 K. Then the systems are cooled down to 293 K within 40 ns and simulated at 293 K to observe for phase separation. The results are shown in Figure S3 (a, b). We have successfully reproduced the phase behavior for these two systems: $(CAG)_{47}$ at concentration 200$\mu$M condenses but $(CAG)_{20}$ at concentration 50$\mu$M does not. More importantly, the result of $(CAG)_{47}$ at concentration 200$\mu$M are obtained in less than 3 days ($\sim$ 60 hours) using 16 CPUs, whereas the original implementation took about 3 months using a GPU for the same system. As shown in Figure S3(c), the measured average concentration of the droplet matches well with the previous reports that the distribution centers around 8 mM.

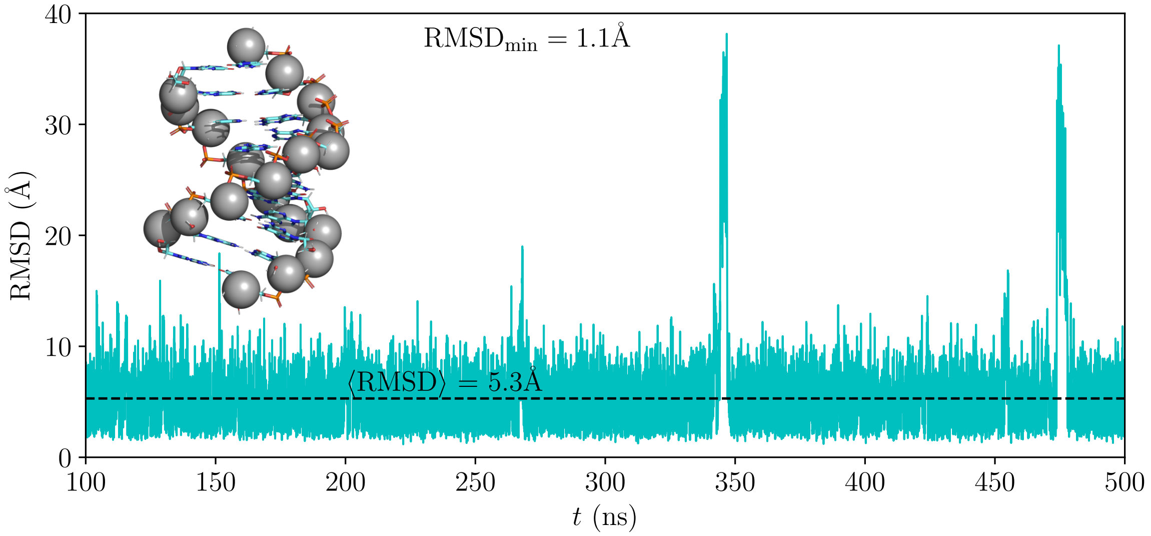


FIG. S2. **Revalidation of the structural property of a single chain that is modified from $(CAG)_2$ duplex (PDB 3NJ6) [3]** by adding an AAAA tetraloop and two terminal A nucleotides. In the canonical simulation at 293 K, the average root-mean-square distance (RMSD) from the crystal structure is about 5.3 Å with a minimum of 1.1Å, which matches the previously reported average of 5.4 Å in the original model implementation. For clarity, the bonds, AAAA tetraloop, and the two terminal A nucleotides are not shown in the included superposition of the minimum RMSD structure.

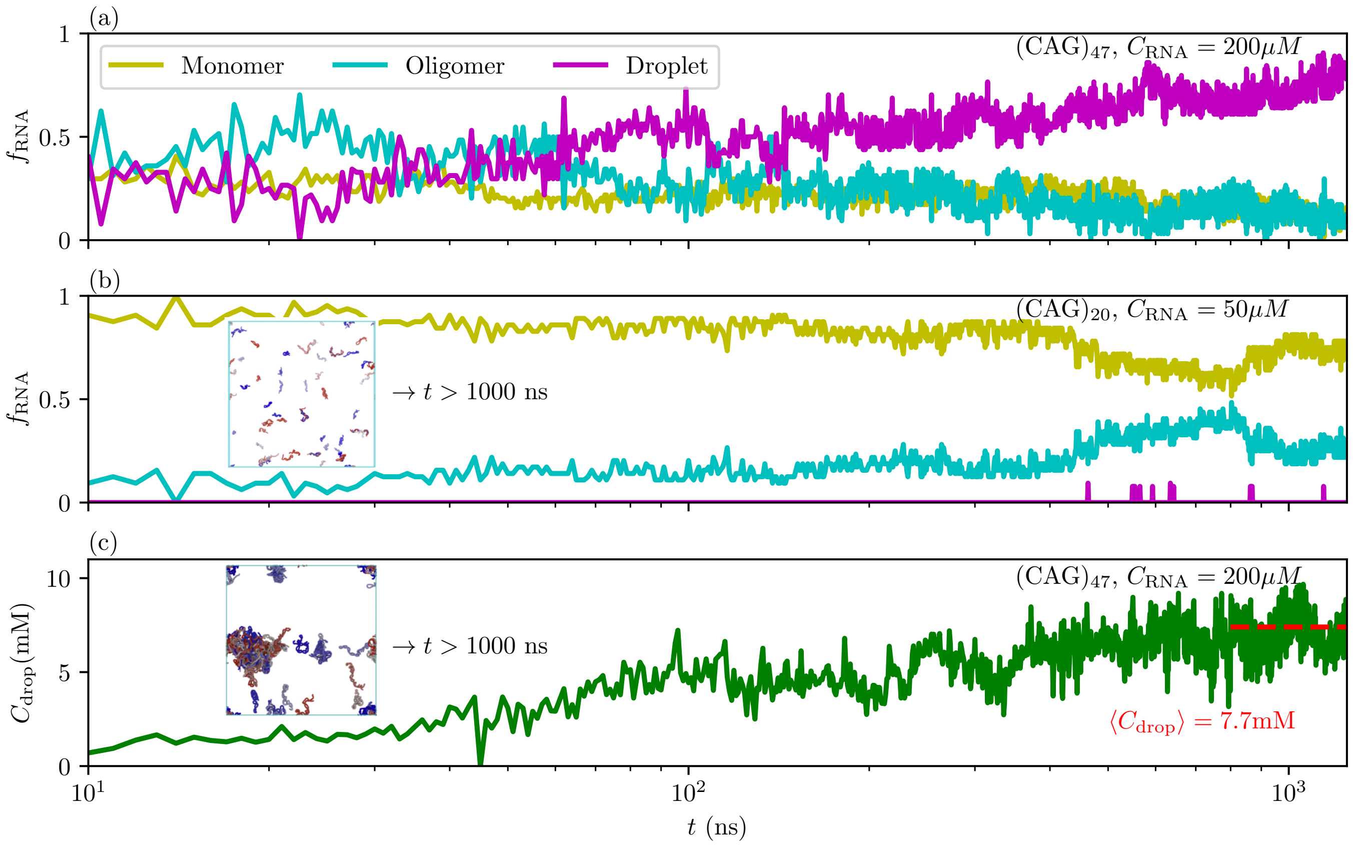


FIG. S3. **Reproduction of condensation results for CAG repeats from previous implementation [4].** **(a)** Time evolution of $(\mathrm{CAG})_{47}$ monomer, oligomer and droplet populations at 200$\mu$M, where monomer is a single chain, oligomer is defined as intermediate clusters with 2–4 chains, and clusters are large clusters with $\geq 5$ chains. We observed consistent dynamics where clusters constituted the predominant composition in $(\mathrm{CAG})_{47}$. **(b)** Time evolution of different structural ensembles in $(\mathrm{CAG})_{20}$ at 50$\mu$M. As in the original simulations, we do not observe phase separation. The less populated oligomers are primarily dimers. **(c)** Concentration of the droplet in $(\mathrm{CAG})_{47}$ from the same simulation as in **(a)**. For the two systems, simulation snapshots ($t > 1000$ ns) are provided in **(b)** and **(c)**.

##### E. Redundancy of RNA trinucleotide repeats

To systematically study the clustering behavior of all trinucleotide repeats, we have also determined the redundancy. As illustrated in Fig. S4, there are 24 non-redundant RNA trinucleotide repeats, and each group of three trinucleotide repeats describes the same chain when repeat numbers are large enough. Since the current model does not consider other interaction types, we have also excluded the homopolymers (poly(rA), poly(rC), poly(rG) and poly(rU)). Therefore, we have simulated 20 non-redundant trinucleotide repeats for this study.

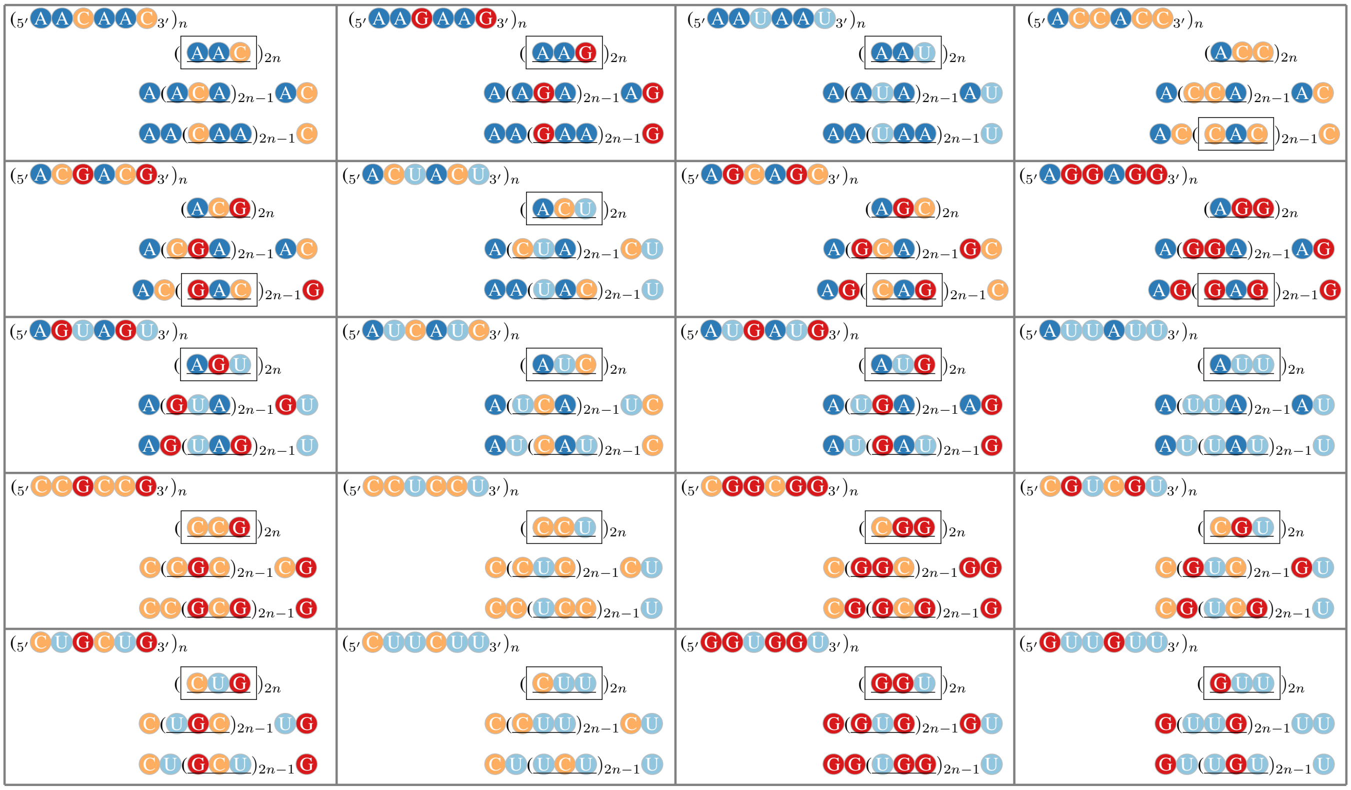

FIG. S4. **Illustration of redundancy in trinucleotide repeats.** Considering the directionality from 5' to 3', there are 64 ($4^3$) combinations. Excluding the 4 homopolymers, there are 60 different trinucleotide sequences. However, as illustrated in each box, when the trinucleotide repeat is long enough (large $n$), each three trinucleotide (underlined) repeat sequences represent the same chain. Therefore, there are only 20 non-redundant trinucleotide repeats. The simulated systems for this paper are shown in square frames.

##### F. Largest clusters and concentrations

Here we also include the time evolution of the size of the largest clusters (Fig. **??**), and their concentrations (Fig. S6).

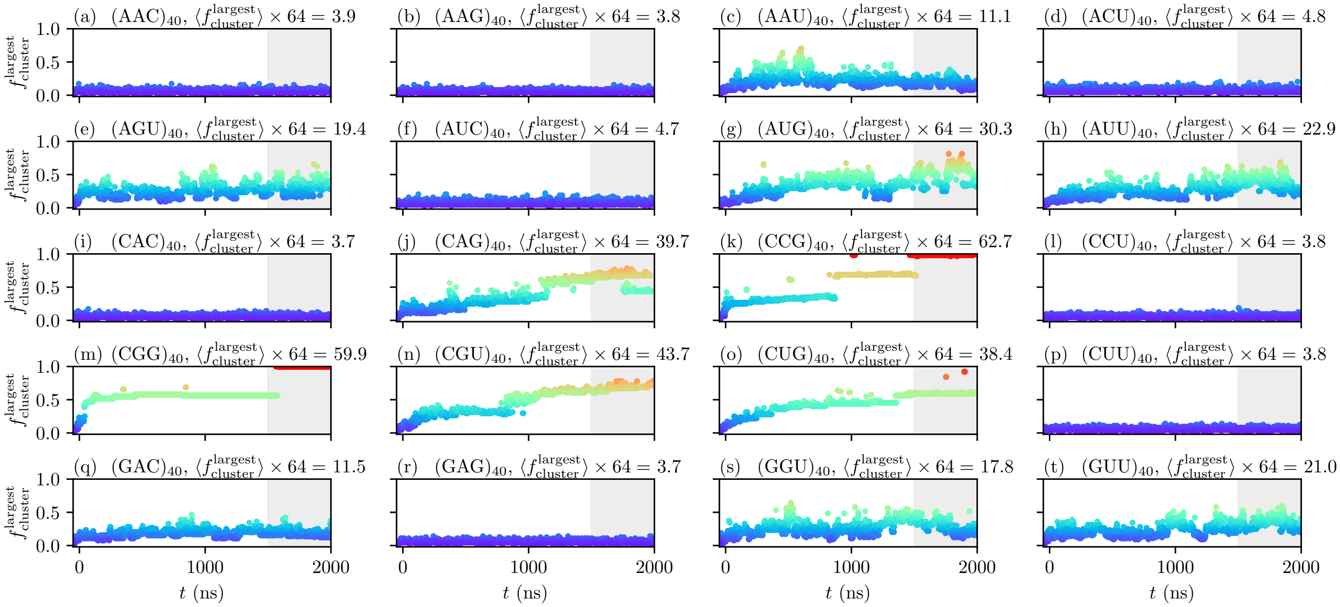


FIG. S5. **Time evolution of the sizes of the largest clusters $f^{\mathrm{largest}}_{\mathrm{cluster}}$ of all 20 non-redundant trinucleotide repeats.** The concentration is calculated from the number of chains in the clusters and the volume of the clusters, the latter being estimated from the gyration tensor of the clusters. Data from the last 500 ns (shaded gray) is used to calculate average values of $f^{\mathrm{largest}}_{\mathrm{cluster}}$ in the system. The data points are colored based on $f^{\mathrm{largest}}_{\mathrm{cluster}}$ values.

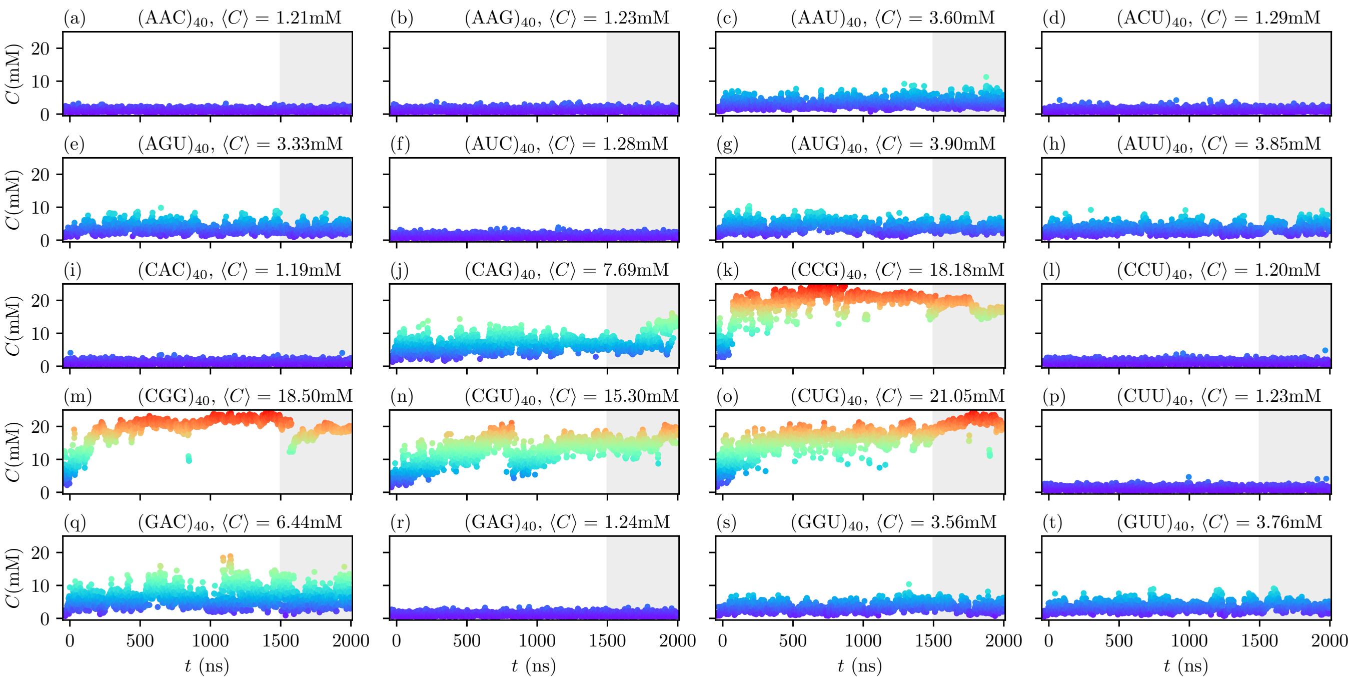


FIG. S6. **Time evolution of the concentrations of the larges clusters $C$ of all 20 non-redundant trinucleotide repeats.** The concentration is calculated from the number of chains in the clusters and the volume of the clusters, the latter being estimated from the gyration tensor of the clusters. Data from the last 500 ns (shaded gray) is used to calculate average values and thehe data points are colored based on $C$ values.

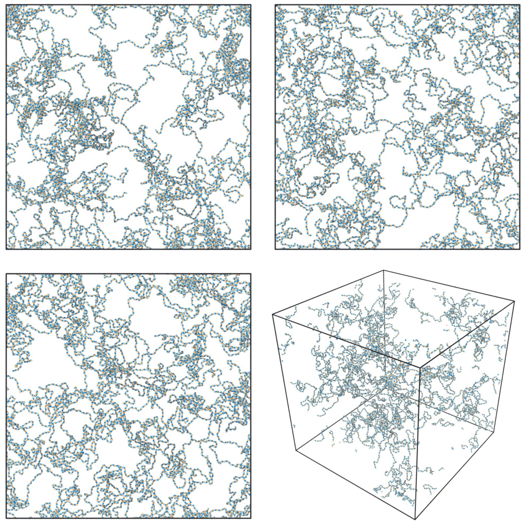

FIG. S7. **Simulation results of (ACU)$_{200}$.** The system is composed of 27 chains and simulated with periodic boundary conditions. The systems are prepared at a similar concentration as (ACU)$_{40}$ at 200 $\mu$M with chains evenly distributed and melted at 373 K. Then the systems are cooled down to 293 K within 40 ns. Finally, the systems are simulated at 293 K for 2 $\mu$s. Consistent with ACU)$_{40}$, within 2 $\mu$s, the system does not phase separate.


[1] C. L. Lawson, H. M. Berman, L. Chen, B. Vallat, and C. L. Zirbel, Nucleic Acids Res. **52**, gkad957 (2023).
[2] W. L. DeLano *et al.*, CCP4 Newsl. Protein Crystallogr **40**, 82 (2002).
[3] A. Kiliszek, R. Kierzek, W. J. Krzyzosiak, and W. Rypniewski, Nucleic Acids Res. **38**, 8370 (2010).
[4] H. T. Nguyen, N. Hori, and D. Thirumalai, Nat. Chem. **14**, 775 (2022).
